# Structure of transmembrane prolyl 4-hydroxylase reveals unique organization of EF and dioxygenase domains

**DOI:** 10.1101/2020.10.25.354423

**Authors:** Matti Myllykoski, Aleksi Sutinen, M. Kristian Koski, Juha P. Kallio, Arne Raasakka, Johanna Myllyharju, Rikkert Wierenga, Peppi Koivunen

## Abstract

Prolyl 4-hydroxylases (P4Hs) catalyze post-translational hydroxylation of peptidyl proline residues. In addition to collagen P4Hs and hypoxia-inducible factor P4Hs, a poorly characterized endoplasmic reticulum (ER)-localized transmembrane prolyl 4-hydroxylase (P4H-TM) is found in animals. P4H-TM variants are associated with the familiar neurological HIDEA syndrome. Here, the 3D structure of the soluble human P4H-TM was solved using X-ray crystallography. The structure revealed an EF-domain with two Ca^2+^-binding motifs inserted to the catalytic domain. A substrate-binding cavity was formed between the EF-domain and the catalytic domain. The active site contained bound Fe^2+^ and N-oxalylglycine. Comparison to homologous structures complexed with peptide substrates showed that the substrate interacting residues and the lid structure that folds over the substrate are conserved in P4H-TM. Differences to homologs were found in the extensive loop structures that surround the substrate-binding cavity and generate a negative surface charge. Ca^2+^-binding affinity of P4H-TM was determined to be within the range of physiological Ca^2+^ concentration in the ER. The proximity of the EF-domain to the active site suggests that Ca^2+^-binding is relevant to the catalytic activity. P4H-TM was found both as a monomer and a dimer in solution, but the monomer-dimer equilibrium was not regulated by Ca^2+^. The solved 3D structure suggests that the HIDEA variants cause loss of P4H-TM function. In conclusion, P4H-TM shares key structural elements with the known P4Hs while possessing a unique property among the 2-oxoglutarate-dependent dioxygenases having an EF-domain and a catalytic activity potentially regulated by Ca^2+^.

## Introduction

Eukaryotic prolyl 4-hydroxylases (P4Hs) are enzymes that catalyze the post-translational hydroxylation of peptidyl-proline residues to 4-hydroxyproline (Fig. 1A). All known P4Hs belong to the same enzyme superfamily of iron and 2OGxoglutarate-dependent dioxygenases (2OGDDs). 2OGDDs are defined by the double-stranded β-helix (DSBH) fold of the catalytic domain, the shared mechanism of the enzymatic reaction and the common cofactors; Fe^2+^, 2OGxoglutarate (2OG), molecular oxygen and vitamin C, which is not a direct cofactor but supports the catalysis (Fig. 1A) (1). Two P4H families with three isoenzymes each have been identified in animals: collagen prolyl 4-hydroxylases (C-P4Hs) 1-3 (2, 3) and the hypoxia-inducible factor (HIF) prolyl 4-hydroxylases (HIF-P4Hs) 1-3, also known as PHDs and EglNs (Fig. 1B) (4, 5). C-P4Hs are α_2_β_2_ heterotetrameric enzymes that are located within the endoplasmic reticulum (ER) and hydroxylate prolines in procollagen α-chains (2, 6). These 4-hydroxyprolines are essential for the stability of the triple helical collagen structure (2, 3, 7). HIF-P4Hs are monomeric enzymes located in cytoplasm and nucleus that specifically hydroxylate the HIFα subunit and mark it for proteasomal degradation via von Hippel-Lindau protein (4, 5). HIF-P4H activity requires high oxygen concentration and these enzymes act as cellular oxygen sensors (8, 9).

**Fig. 1.**
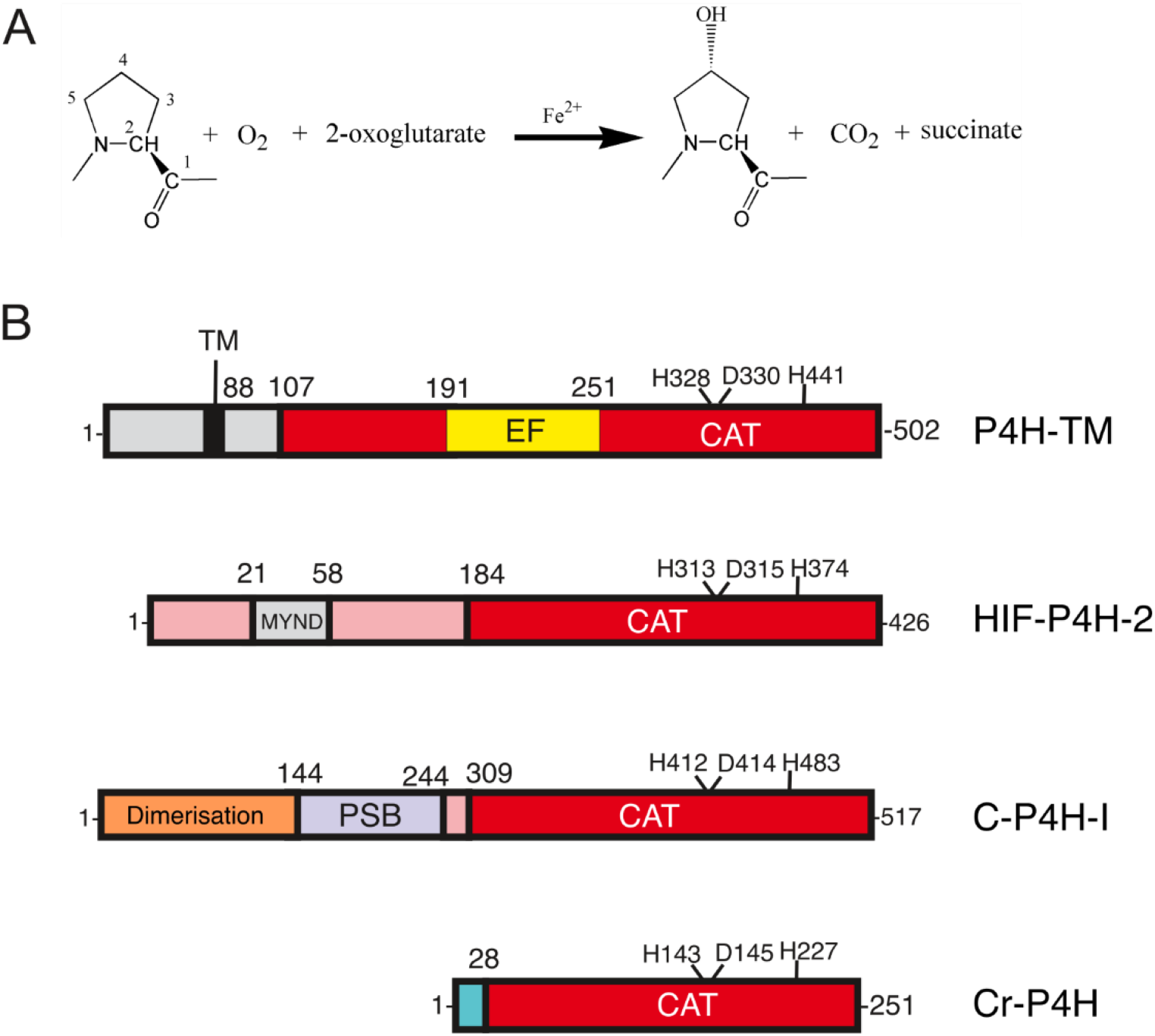
The reaction catalyzed by P4Hs and the domain assignments of selected P4Hs. A) All P4Hs share the same reaction mechanism and cofactors. B) P4H-TM, HIF-P4H-2, the α-subunit of human C-P4H-I and *C. reinhardtii* P4H (Cr-P4H) share the DSBH fold of the catalytic domain (CAT, red) and the catalytic residues (which are indicated), but they have unrelated N-terminal regions. P4H-TM has a cytosolic N-terminal region, a transmembrane helix and an EF-domain inserted into the catalytic domain. The structure of the N-terminal region of HIF-P4H-2 is not known but it is predicted to contain a MYND-type zinc finger (15). The N-terminal half of the α-subunit of human C-P4H is well characterized and contains a dimerization domain followed by a peptide-substrate-binding (PSB) domain (6). Structural information is missing for the C-P4H catalytic domain and for the linker region between the PSB and the catalytic domain. The functional C-P4H enzyme is an α_2_β_2_ heterotetrameric complex between the catalytic α-subunit and the β-subunit/protein disulfide isomerase (not shown). Cr-P4H represents the simplest type of P4H that is lacking an extended N-terminus (16).

Transmembrane prolyl 4-hydroxylase (P4H-TM) is considered to be the fourth HIF-P4H (10, 11). It is located at the ER membrane with the catalytic domain inside the ER lumen (11). P4H-TM sequence resembles more closely the C-P4Hs than the HIF-P4Hs (Fig. 2), but instead of procollagen, it was found to hydroxylate HIFα *in vitro* and to downregulate HIFα in cell culture (10, 11). Two putative EF-hand motifs were detected in the P4H-TM sequence N-terminal to the catalytic domain (Fig. 1B, Fig. 2) (10). The Ca^2+^-binding EF-hand motifs that were first identified in parvalbumin by Kretsinger (12), are around 30-residue long helix-loop-helix structures that usually occur in pairs. Ca^2+^-binding by seven oxygen groups within the EF-hand loop region modulates the relative orientation of the two helices (13, 14). EF-hand containing proteins can function as calcium sensors, generating biochemical responses to changes in cellular calcium concentration, or as calcium buffers, binding free cellular calcium to modulate cellular signaling (13).

**Fig. 2.**
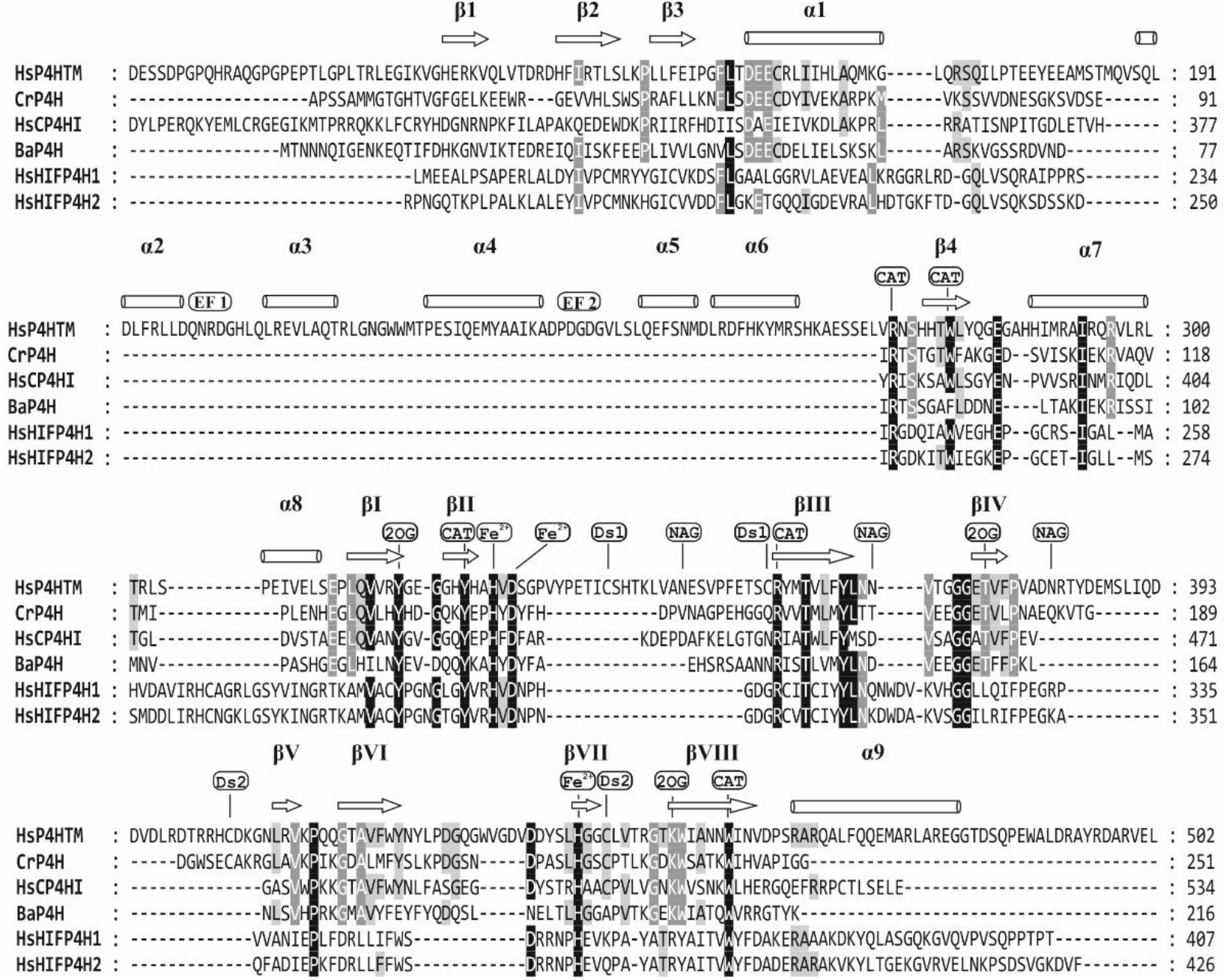
Structure-based sequence alignment of the catalytic domains of selected P4Hs. The alignment includes human P4H-TM (HsP4HTM), *C. reinhardtii* P4H (CrP4H), human C-P4H isoform I (HsCP4HI), *Bacillus anthracis* P4H (BaP4H) and the isoforms 1 and 2 of human HIF-P4Hs (HsHIFP4H1 and HsHIFP4H2, respectively). The presented sequence of P4H-TM starts with Asp88. The secondary structure elements indicated above the sequences are based on the crystal structure of the P4H-TM determined in this study. The β-strands that form the DSBH core are labelled with Roman numerals. The conserved residues are highlighted, and the residues involved in iron chelation (Fe^2+^: His328, Asp330 and His441), 2OG binding (2OG: Tyr319, Thr375 and Lys451) and P4H activity (CAT: Arg273, Trp279, Tyr325, Arg358 and Trp457) are labeled. Disulfides (Ds) and glycosylated (NAG) residues are also labeled. A distinct feature in P4H-TM compared to the other P4Hs is the EF-domain between residues Gln190 and Asn251 that contains two EF-hands (EF1 & EF2).

P4H-TM is highly expressed in brain and eye and moderately expressed in skeletal muscle, lung, heart, adrenal gland and kidney (11, 17, 18). Expression has also been reported in prostate, testis and thyroid (11). Morpholino knockout of P4H-TM in zebrafish embryos resulted in basement membrane defects, impaired eye development and compromised kidney function (17). In mice, P4H-TM is involved in regulation of erythropoietin levels and erythrocytosis (19). *P4h-tm*^−/−^ mice develop early-onset aging-associated retinal and renal dysfunction (18), and their behavioural phenotype is characterized by hyperactivity and a dramatic reduction of despair response (20).

Variants of human *P4HTM* have been linked to a severe disability, the HIDEA syndrome, characterized by intellectual disability, hypotonia, eye abnormalities, hypoventilation, obstructive and central sleep apnea and dysautonomia (21, 22). Exome sequencing revealed five different homozygous or compound heterozygous pathogenic biallelic *P4HTM* variants in patients from five families from across the world (21, 22). Two of the variants lead to premature stop codons and the remaining three resulted in insoluble protein products, suggesting the disease is linked to the loss of P4H-TM function (22).

Although previous results show that P4H-TM hydroxylates HIF1α and it has been considered to be the fourth HIF-P4H, the catalytic activity was only detected towards the 200-residue oxygen-dependent degradation domain (ODDD) of HIF1α, not towards shorter peptides harboring the prolines whose hydroxylated forms are recognized by von Hippel Lindau protein (11). Furthermore, P4H-TM also hydroxylated to a small extent HIF1α ODDD in which the HIF-P4H targeted prolines were mutated to alanines. In addition, the behavioral phenotype of the P4H-TM knockout mice is very different from any other HIF-P4H knockouts (20), and the symptoms of the HIDEA syndrome patients have not been reported to be linked to HIF-P4H deficiency (21, 22). P4H-TM contains a putative EF-domain with calcium-binding EF-hand motifs not found in any other characterized 2OGDD superfamily enzyme. The significance of this domain for P4H-TM function, the potential connection of calcium storing/sensing to HIF1α regulation, and the relevance of P4H-TM localization inside the ER are not known. In order to shed light on the function of P4H-TM, we solved the crystal and solution structures of the protein, analyzed the structures and used them to predict the effect of the HIDEA variants on the structure and function within the cellular context and in relation to the calcium concentration.

## Results

### Overall structure of human P4H-TM

The structure of the soluble part of human P4H-TM was solved with X-ray crystallography. The crystallized construct consists of residues 88-502 of P4H-TM and an N-terminal His-tag while it lacks the short cytoplasmic region and the transmembrane helix (Fig. 1). The crystal structure is composed of two well-defined domains: the catalytic domain with the DSBH fold and the EF-domain inserted into the middle of the catalytic domain (Fig. 3). Overall, the structure contains twelve β-strands and nine α-helices. The conserved DSBH core is composed of eight antiparallel β-strands that are divided into two sheets that fold against each other. The core strands are conventionally numbered with roman numerals from I to VIII (Fig. 3A). Two of the core strands of the DSBH minor sheet are heavily disrupted in P4H-TM and do not observe the β-sheet geometry, but the nomenclature is preserved for clarity. The two P4H-TM molecules of the asymmetric unit occupy very similar conformations (RMSD 0.42 Å for all Cα-atoms), and only the chain A is discussed unless otherwise stated.

**Fig. 3.**
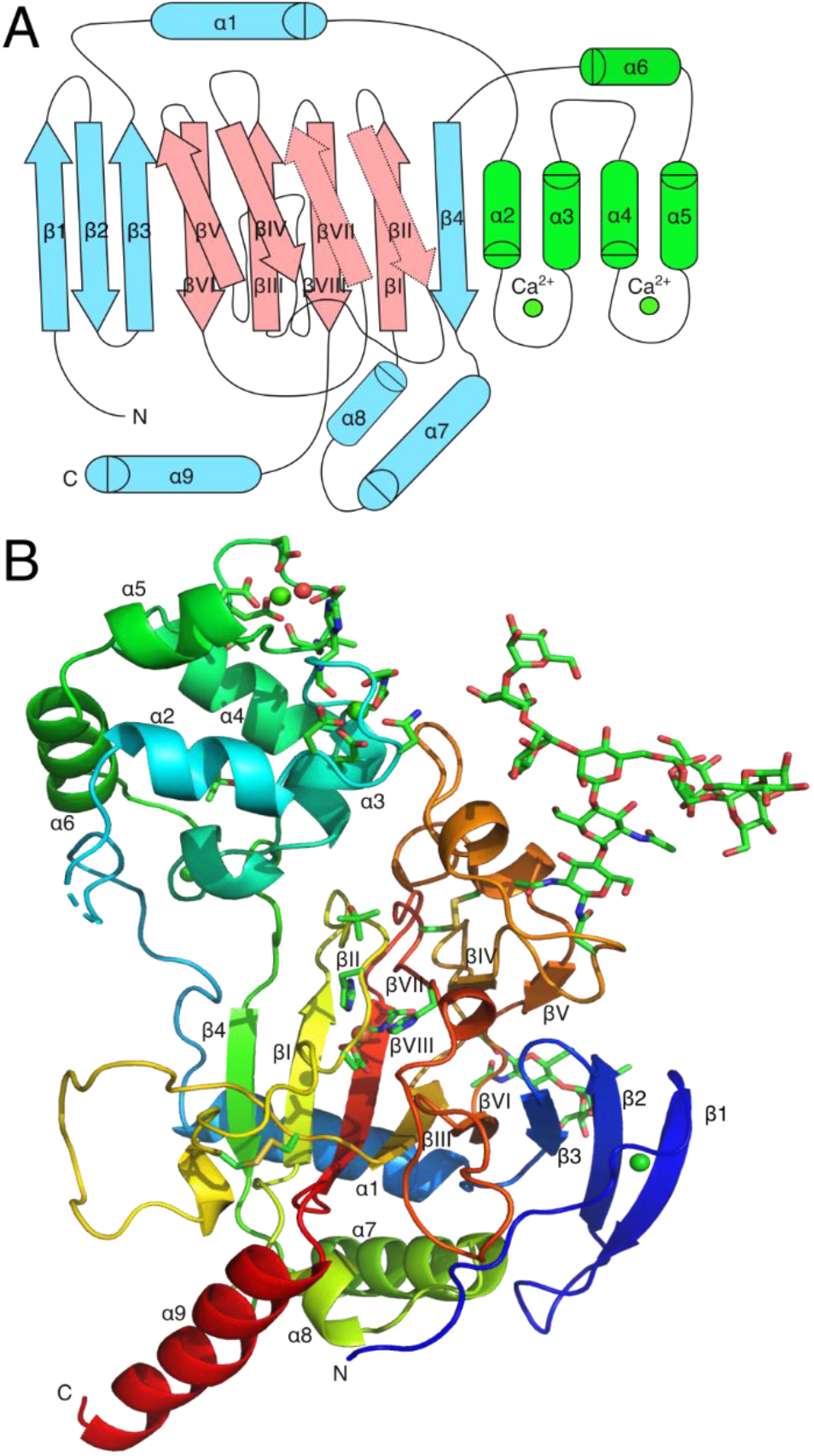
P4H-TM structure. The structure is presented as A) a topology chart where the EF-domain is shown in green, the DSBH core in red and the rest of the protein in cyan and B) a rainbow-colored cartoon model.

The first visible P4H-TM residues are Thr107 and Leu108 in chains A and B, respectively. Including the 6xHis tag, there are 25 disordered residues in the N-terminus that are not visible in the electron density. The visible N-terminus is positioned along the protein surface between helices α7 and α8 and the βVI-βVII loop without defined secondary structure. Starting at Gly120, the protein forms three consecutive antiparallel β-strands (β1-3) that extend the DSBH major sheet. After β3, the first helix α1 is positioned behind the major sheet. α1 is followed by a long and partially disordered loop α1-α2 which reaches the EF-domain. The first of the two EF-hands is formed by α2, a calcium-binding loop α2-α3 and α3. α3 ends abruptly after six residues and is followed by a 3_10_ helix and the α3-α4 loop between the two EF-hands. The second EF-hand is formed by helices α4 and α5 and the calcium-binding loop α4-α5 in between them. The six-residue α5 is followed, after a turn, by the longer α6. The loop α6-β4 runs antiparallel to the α1-α2 loop back towards the catalytic domain and forms a two-residue β-like extension to the DSBH minor sheet. β4 extends the DSBH major sheet to the opposite direction to the strands 1-3. It is followed by two helices, α7 and short α8, that run antiparallel to each other and lead into the first β strand of the DSBH core. βI is located between β4 and βVIII in the DSBH major sheet. βII folds over βI, but its conformation is disrupted by the iron-binding residues and the disulfide in the neighboring βVII strand. βII leads to an extended 30-residue loop βII-βIII, which contains an internal disulfide between Cys340 and Cys357. The disulfide bridge appears to tether the middle part of the loop to the beginning of βIII. βIII is part of the major sheet between βVI and βVIII. Subsequent βIV folds over it, positioned between βV and βVII, and leads to a 30-residue βIV-βV loop that harbors two 3_10_ helices and forms several contacts with the EF-domain. A disulfide is formed between Cys404 near the end of the βIV-βV loop and Cys444 at the end of βVII. This linkage seems to function as an anchor to determine the position of this large loop. After the loop, βV forms one end of the minor sheet and is positioned antiparallel to the neighboring βIV. A short turn after βV leads to βVI positioned between β3 and βIII. The following loop, βVI-βVII twists around itself in an extended hairpin-like structure and leads to βVII after a short 3_10_ helix. βVII completes the minor sheet between βII and βIV and is followed by the final major sheet strand βVIII. A short, coiled region after βVIII leads to the terminal helix α9. The last visible residue in the structure is Gly481 at the end of α9, only 25 Å away from the visible N-terminus. The last 21 residues of P4H-TM are not visible in the electron density.

Prediction of glycosylation sites within P4H-TM resulted in three potential sites of which two, Asn368 and Asn382, reached the threshold while the third site, Asn348, was just below it. In the structure, Asn368 and Asn382 were found to be glycosylated, while Asn348 is located in a part of the βII-βIII loop where the electron density for the side chain was not well defined. Asn368 is in the short βIII-βIV loop and two N-acetylglucosamine residues could be modelled to be attached to it with clear electron density. Eight mannose residues were modelled to the Asn382-linked glycan in the βIV-βV in addition to the two N-acetylglucosamines.

A suggested alternative isoform 3 of P4H-TM (Uniprot: Q9NXG6-3) to isoform 1 (Uniprot: Q9NXG6-1) contains a 61-residue insertion after Arg358 and appears in some sequence databases as the canonical form of the enzyme (Fig. S1). We expressed isoform 3 with a similar insect cell expression vector used for the isoform 1 lacking the cytosolic part and the transmembrane domain, but did not obtain soluble protein for characterization (data not shown). P4H-TM structure indicates that an insertion at this position would either displace Arg358 and triple the length of the βII-βIII loop or alternatively completely displace the βIII and βIV strands.

### EF-hand and Ca^2+^-binding

P4H-TM structure contains an EF-domain with two EF-hands between residues 190 and 251 (Fig. 4A). Calcium is coordinated in a pentagonal bipyramid conformation by 7 oxygen ligands from residues listed in Table 1. The initial Ca^2+^-binding residue Asp198 in the first EF-hand emerges directly from α2. The -Y position of the first EF-hand is provided by the main chain carbonyl of His204 while Glu209 links α3 to the Ca^2+^ with bidentate binding. Unusually, the initial Ca^2+^-binding residue of the second EF-hand, Asp237, does not emerge from α4 directly, but after a linker residue. This displacement could influence the relative movement by α4 and α5 in the event of Ca^2+^ dissociation. The coordinating oxygen in the -X position of the second EF-hand is provided by a water molecule in chain A. No electron density for a corresponding water molecule was visible in chain B. The main chain carbonyl oxygen of Val243 provides ligand position −Y and Glu248 from α5 binds Ca^2+^ in a bidentate manner.

**Table 1.** EF-domain Ca^2+^-coordinating residues of P4H-TM.

| **Position** | **EF 1** | **EF 2** |
| --- | --- | --- |
| **X** | Asp198 | Asp237 |
| **Y** | Asn200 | Asp239 |
| **Z** | Asp202 | Asp241 |
| **-X** | Gln206 | Water 33 |
| **-Y** | His204 | Val243 |
| **-Z** | Glu209 | Glu248 |

**Fig. 4.**
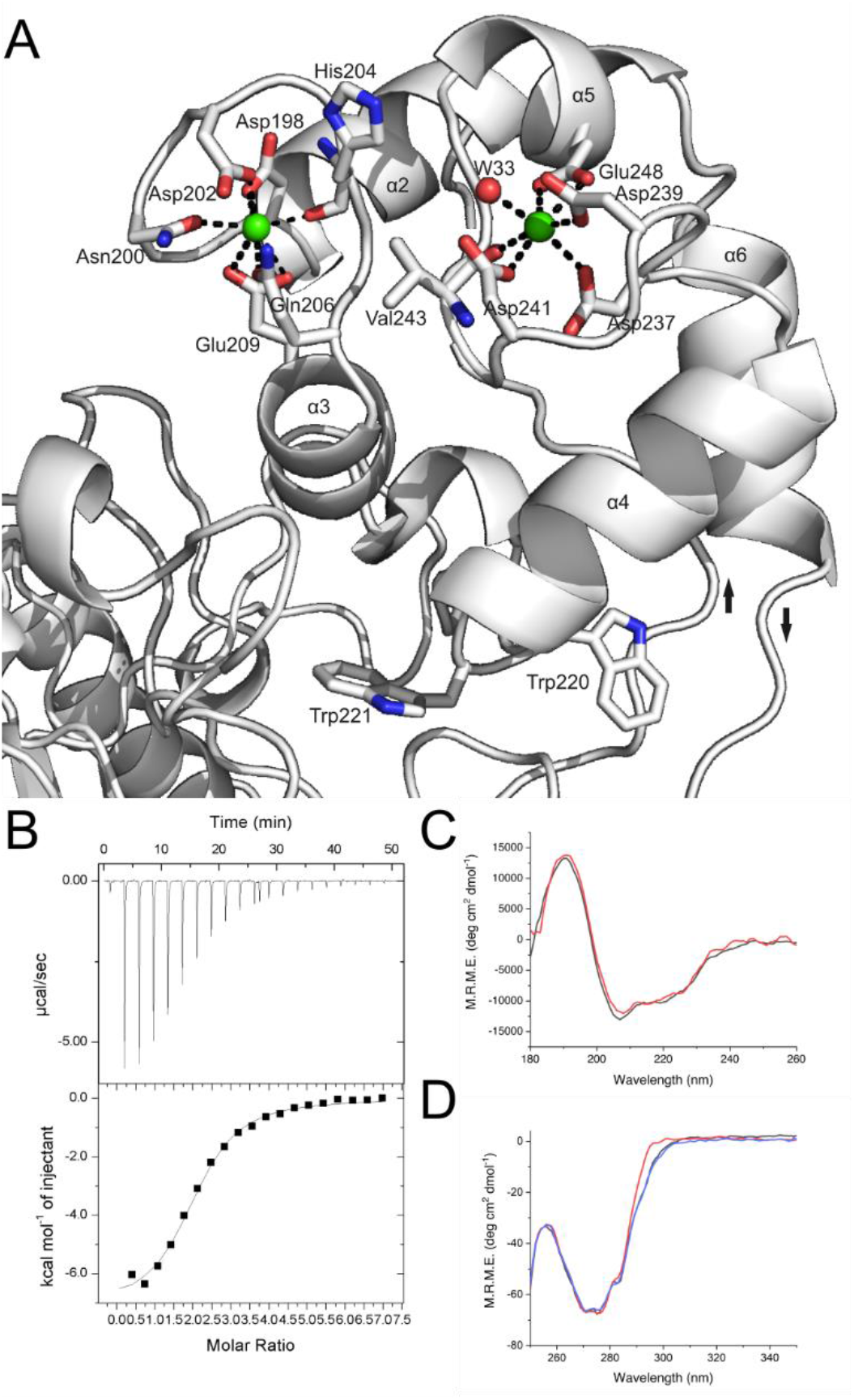
Calcium binding by the EF-domain of P4H-TM. A) Structural representation of the EF-domain of P4H-TM. Calcium interacting residues and secondary structure elements are labeled. Calcium ions are shown in green. The incoming α1-α2 and outgoing α6-β4 loops are marked with arrows that indicate the direction of the polypeptide chain. B) Isothermal titration calorimetry plot from an injection of CaCl_2_ to P4H-TM. C) Far-UV synchrotron radiation circular dichroism (CD) of P4H-TM between wavelengths 180 nm and 260 nm without (black) and with CaCl_2_ (red). D) Near-UV CD of P4H-TM between wavelengths 250 nm and 350 nm without metals (black), with CaCl_2_ (red) and with MgCl_2_ (blue).

The overall Ca^2+^-binding affinity of P4H-TM was measured using isothermal titration calorimetry (ITC) (Table 2) (Fig. 4B). The modeling used assumes that the two binding sites are identical and independent. The obtained data fit well to this model and the stoichiometry is accurate. The results suggest that calcium binding to P4H-TM is enthalpy-driven as the negative ΔH term is nearly nine times larger than the slightly positive entropy term −TΔS (Table 2).

**Table 2.** Isothermal titration calorimetry results of P4H-TM.

|  |  |
| --- | --- |
| **N** | $2.02 \pm 0.039$ sites |
| **K** | $42700 \pm 5030$ M<sup>-1</sup> * |
| **ΔH** | $-7142 \pm 186.6$ cal mol<sup>-1</sup> |
| **ΔS** | $-2.77$ cal mol<sup>-1</sup> deg<sup>-1</sup> |

N, molar ratio of Ca<sup>2+</sup> binding; K, Ca<sup>2+</sup> binding constant; ΔH, enthalpy change; ΔS, entropy of binding.
\*Corresponds to $K_d$ 23 μM.

Circular dichroism (CD) measurements were used to clarify the structural changes that occur during calcium binding to P4H-TM. Far-UV synchrotron radiation CD (SRCD) measurements did not show a notable change in the secondary structure composition of P4H-TM with the addition of calcium (Fig. 4C). In contrast, the near-UV CD revealed a large shift in the CD signal in the tryptophan region between wavelengths 285 nm and 305 nm when calcium was added to metal-free P4H-TM (Fig. 4D). No shift was induced when magnesium was added to metal-free P4H-TM (Fig. 4D). This signal probably arises when the calcium-dependent conformational change shifts the positions of Trp220 and Trp221 located in the α4-α5 loop between the two EF-hands (Fig. 4A). Trp220 interacts with the flexible α1-α2 loop and forms CH-π interaction with Pro175. Trp221 interacts with the βIV-βV loop and is stacked with His326.

### Active site

The active sites of 2OGDD family enzymes are remarkable in that the binding of iron and 2OG is highly conserved, but there is a large variation in the binding modes of the hydroxylatable substrates. The P4H-TM substrate-binding cavity is located in between the EF-domain and the catalytic domain. The active site is located at the catalytic domain side of the substrate-binding cavity and contains bound iron and the 2OG analog N-oxalylglycine (NOG). The iron is coordinated by His328, Asp330, His441 and the oxygen atoms from C-1 carboxylate and C-2 carbonyl of NOG (Fig. 5, Fig. S2). NOG C-1 carboxylate group is shifted above the plane formed by Asp330, His441 and the NOG C-2 carbonyl group, that is unlike in a typical 2OGDD active site where the iron-coordinating residues follow octahedral geometry. Additionally, there is no water molecule positioned trans to His328. NOG is coordinated at the C-5 carboxyl group by Tyr319, Thr375 and Lys451 and at the C-1 carboxyl by Asn455 (Fig. S2). The interaction between NOG and Asn455 may contribute to the disrupted iron coordination geometry. The P4H-TM iron-binding residues His328, Asp330 and His441 are conserved in other P4Hs (Fig. 2). Of the NOG-binding residues only Tyr319 is fully conserved (Fig. 2). Lys451 and Thr375 are conserved, except in HIF-P4Hs where the lysine is replaced by an arginine and the threonine with a leucine (Fig. 2). Further, in HIF-P4Hs the conserved tyrosine corresponding to Tyr365 in P4H-TM interacts with the co-substrate, while in P4H-TM Tyr365 forms a hydrogen bond to the co-substrate interacting Thr375. Asn455, although conserved in C-P4H-I, is replaced in most P4Hs by a threonine that does not form similar interaction with the co-substrate (Fig. 2). NOG-binding in P4H-TM is also altered, compared to homologs, by Gly443, which in homologs usually has a side chain that restricts the 2OG/NOG binding site (Fig. 2). Iron and 2OG/NOG coordinating residues are well-conserved in P4H-TM orthologs (Fig. S3).

**Fig. 5.**
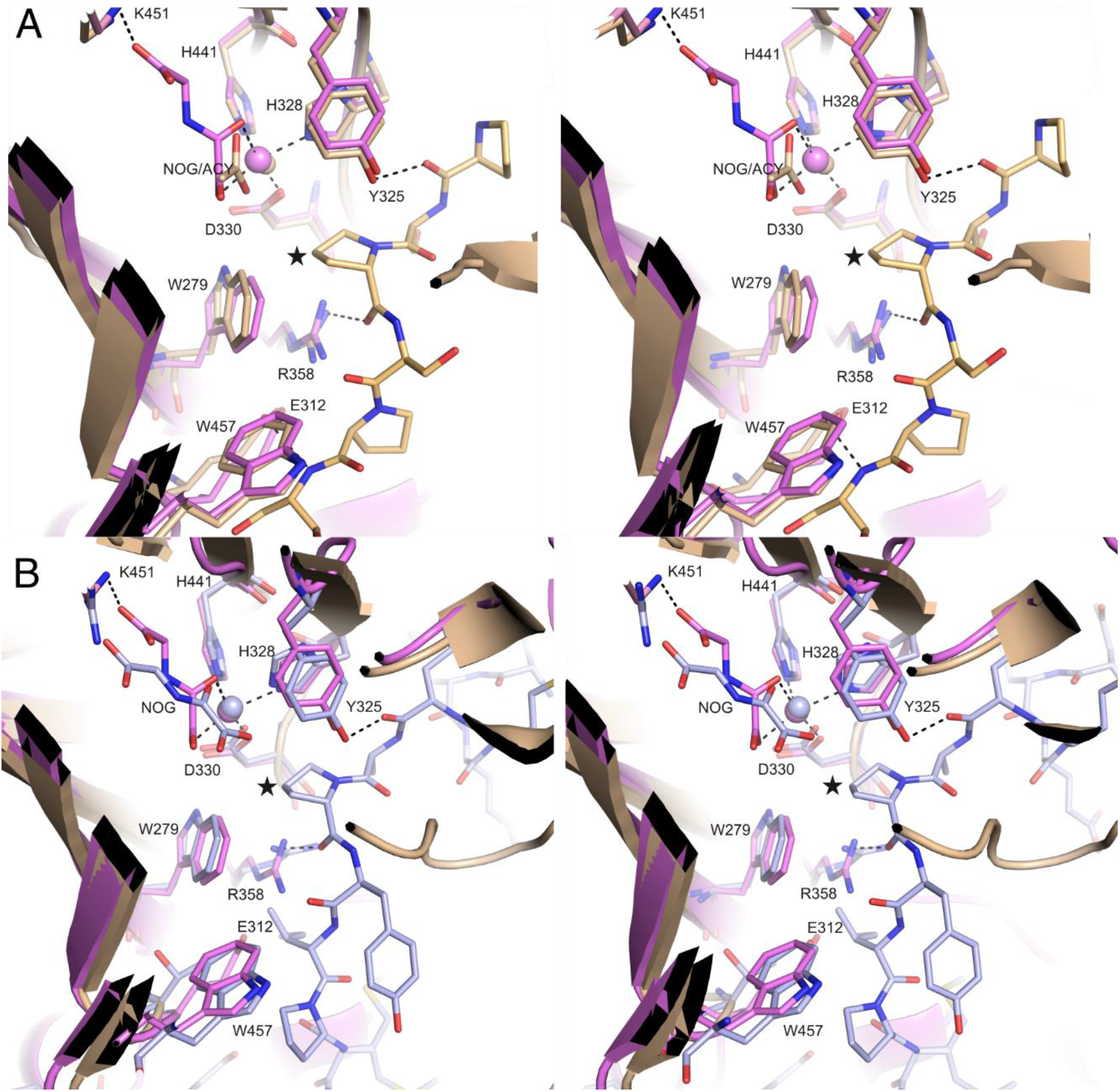
Comparison of the P4H-TM active site to homologous structures. Stereo figures of the structural organization of the P4H-TM active site (violet) compared to the peptide-bound homologous structures of A) *C. reinhardtii* P4H (light brown) and B) HIF-P4H-2 (light blue) active sites displaying the iron and 2-oxoglutarate analogue N-oxalyl glycine (NOG) coordinating residues and the P4H activity-linked residues. Highlighted P4H-TM residues are numbered and the hydroxylated prolines of the substrate peptides are marked with stars.

The residues Arg273, Trp279, Glu312, Tyr325, Arg358 and Trp457 near the P4H-TM active site are conserved in other P4Hs and function in substrate binding or catalysis. The function of these residues can be predicted based on the substrate-peptide containing P4H structures of *C. reinhardtii* P4H (Cr-P4H) and HIF-P4H-2 (Fig. 5). In these homolog structures, Arg273 interacts with the first residue of the lid structure and with the substrate peptide via a water molecule. Trp279 forms stacking interaction with a peptide bond of the substrate peptide in the Cr-P4H structure. Glu312 is not conserved in HIF-P4H-2 but interacts with the substrate peptide and Arg358 in the Cr-P4H structure. Tyr325 is stacked between Arg273 and His328, forms a hydrogen bond to the backbone carbonyl of the substrate peptide residue at position −2 to the hydroxylated proline and is in intimate proximity of the hydroxylated proline. Arg358 directly interact with the backbone carbonyls of the hydroxylated proline and Asp330. Trp457 forms a stacking interaction with Arg358 and is hydrogen bonded with the carboxylate group of Asp330. These residues are also completely conserved in P4H-TM orthologs (Fig. S3). The presence of conserved residues linked to P4H activity indicates that the central aspects of P4H function and substrate binding are conserved in P4H-TM.

Four loop structures surround the P4H-TM active site (Fig. 6A). The loop α1-α2 leads from the catalytic domain to the EF-domain next to the active site. The βII-βIII loop with an internal disulfide borders the active site cavity on one side. The βIV-βV loop forms interactions with the EF-domain, and together with the loop α3-α4 from the EF-domain makes the substrate-binding cavity of P4H-TM longer and narrower compared to homologous enzymes. The βVI-βVII loop extends the length of the cavity next to βII-βIII and opposite to βIV-βV. The sequences of the P4H-TM loops βII-βIII, βIV-βV and βVI-βVII are almost completely conserved among vertebrates and any substrate-interacting residues located there are likely to be preserved (Fig. S3). All loops are present also in Cr-P4H but they are shorter and the sequence conservation to P4H-TM is very limited. In addition, the βII-βIII loop occupies the active site in the Cr-P4H structure with zinc and pyridine 2,4-dicarboxylate but without the peptide substrate (Fig. 6C) (16). In HIF-P4H-2 these loops are practically absent, and the position of βII-βIII is occupied by the C-terminal helix that also interacts with the substrate peptide. The cysteines 340 and 357 that form a disulfide in the βII-βIII loop, are not conserved elsewhere (Fig. 2). On the other hand, Cys444 is conserved in Cr-P4H and C-P4H-I, but Cys404 is conserved only in Cr-P4H, suggesting that the corresponding disulfide, if present, is formed differently in C-P4H-I (Fig. 2). The disulfide-forming cysteines are completely conserved in P4H-TM orthologs, but some invertebrate proteins have additional cysteines in the βII-βIII loop (Fig. S3).

**Fig. 6.**
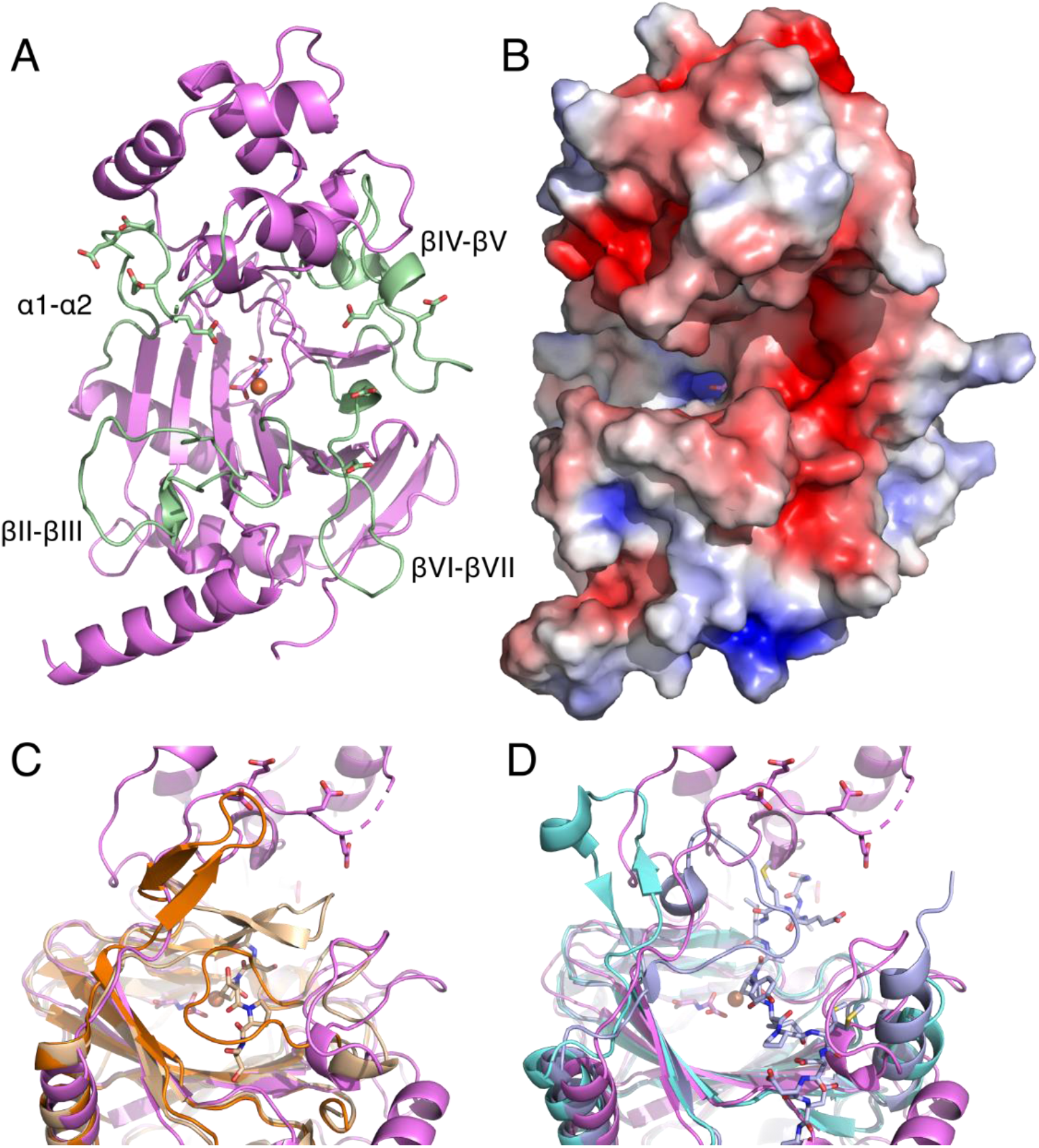
Comparison of the peptide substrate binding cavity of P4H-TM, *C. reinhardtii* P4H (Cr-P4H) and HIF-P4H-2. A) The loops surrounding the P4H-TM (violet) active site are labeled and highlighted in light green. The side chains of the acidic residues are also shown. B) The electrostatic surface around the active site shows negatively charged surface brought about by the acidic residues. Cartoon in A) and surface in B) are in the same orientation. The opened and closed lid structures of the active site of C) Cr-P4H and D) HIF-P4H-2 overlaid with the P4H-TM active site with the open conformation. Cr-P4H open (PDB id 2jig) and closed (PDB id 3gze) conformation structures are shown in orange and light brown, respectively. HIF-P4H-2 open (PDB id 2g19) and closed (PDB id 3hqr) conformation structures are shown in cyan and light blue, respectively.

Electrostatic surface calculation shows that the substrate binding cavity is lined with negatively charged residues resulting in an overall negative charge concentrated on two positions (Fig. 6B). The first is at the opening of the cavity where the most prominent acidic residues are Asp386 and Glu387 from βIV-βV loop and Asp434 and Asp437 from βVI-βVII loop. The second is at the other end of the cavity and is composed of glutamates 177, 178, 180 and 181 from the α1-α2 loop. The corresponding residues in Cr-P4H and HIF-P4H-2 are part of a lid structure on top of the substrate peptide (Figs. 6C and 6D), suggesting that these glutamates may be involved in a similar role. The acidic residues are nearly always conserved in vertebrate P4H-TM sequences as either aspartate or glutamate residues (Fig. S3). In comparison, Cr-P4H active site also contains negatively charged residues but they are much less prominent than in P4H-TM (Fig. S4). On the other hand, in HIF-P4H-2 the HIFα-binding site is largely positively charged (Fig. S4). The differences in the P4H-TM active site compared to homologs outside the immediate core suggest a different substrate peptide or a different substrate binding mode than in the other enzymes.

Cr-P4H and HIF-P4H-2 peptide-bound structures have a lid structure folded over the substrate peptide (Figs. 6C and 6D). In peptide-free structures this region is either disordered or folded away from the active site (Figs. 6C and 6D). In P4H-TM this lid structure seems to be conserved and is formed by the partially disordered α1-α2 loop.

### Solution structure of P4H-TM and the impact of calcium

The calcium-bound P4H-TM crystal structure does not adequately clarify the role of Ca^2+^ in P4H-TM function. However, P4H-TM did not crystallize without Ca^2+^. To overcome this, the conformation change caused by calcium binding was modelled using the structure of calmodulin N-terminal EF-hand pair without calcium. The Ca^2+^loss in calmodulin leads to a relative shift in the positions of the EF-hand helices. P4H-TM without calcium was modelled in three ways with the EF-domain helices fixed-in-place at three different positions (Figs. 7A-C). The model where α3 was fixed resulted in the decrease of the overall length of the protein as the result of Ca^2+^ loss (Fig. 7B). This alternative would mostly conserve the substrate-binding cavity and the interactions between the EF-domain and the βIV-βV loop. In the two other models, where either α2 position was fixed (Fig. 7A), or α5 was fixed to extend α6 (Fig. 7C), the whole EF-domain moved away from the catalytic domain and increased both the size of the substrate-binding cavity and the overall length of the protein. These models would likely disrupt the interactions between the EF-domain and the βIV-βV loop, unless the loop is capable of substantial elongation.

**Fig. 7.**
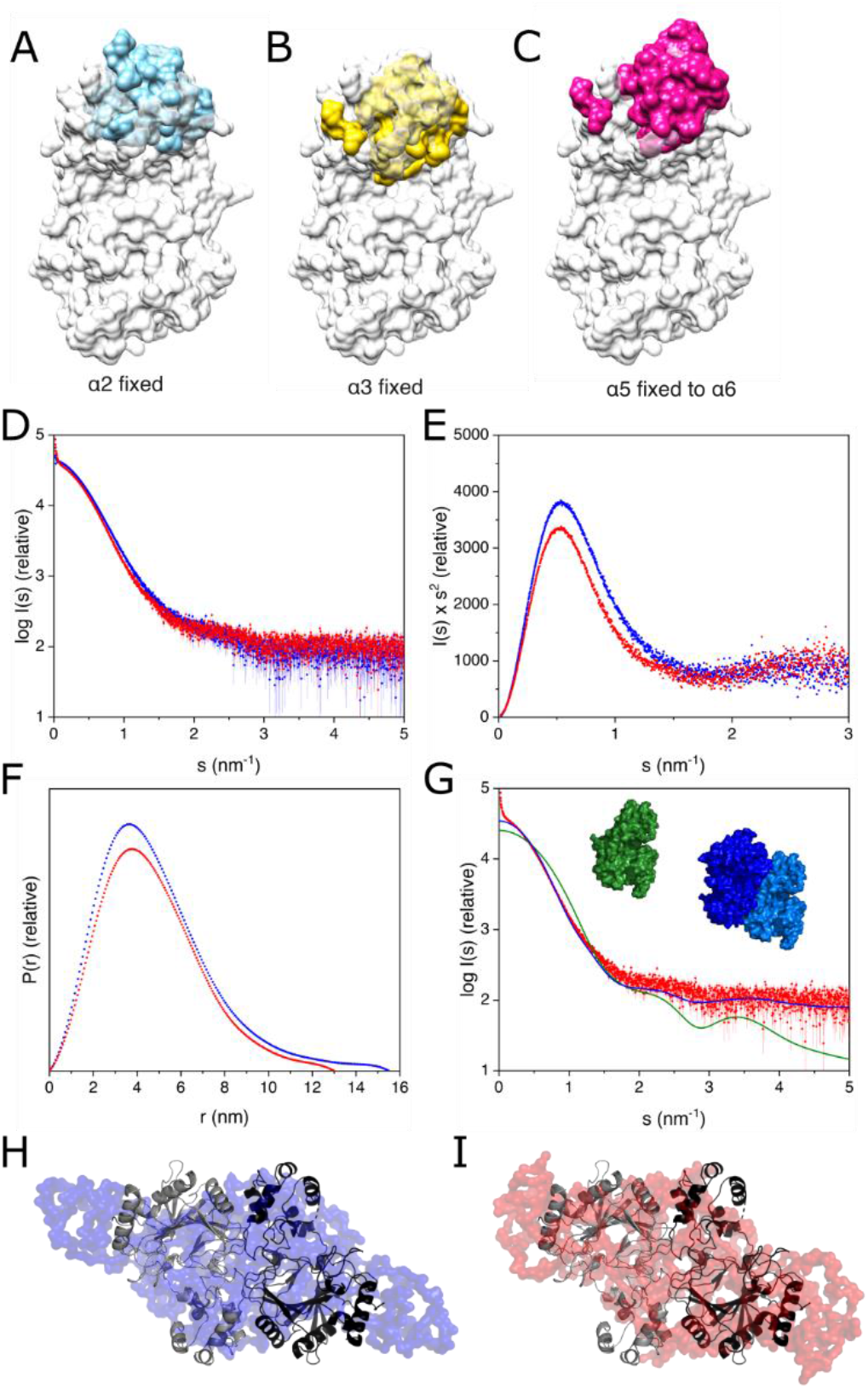
Modelling of the effect of Ca^2+^ loss to crystal and solution structures of P4H-TM. A-C) P4H-TM monomer structure where EF-domain is morphed to resemble the orientation of Ca^2+^-unbound calmodulin in conformations where A) α2 or B) α3 were fixed in place or C) where α5 was fixed to extend α6. The crystal structure is shown in white and the unbound EF-domain models are shown in A) cyan, B) yellow and C) magenta. D-G) SAXS results from P4H-TM in solution with Ca^2+^ (red) and without Ca^2+^(blue). D) Scattering data with and without Ca^2+^ overlaid. E) Kratky plot indicating no difference in the overall folding with and without Ca^2+^. F) Distance distribution of the two forms in solution. Maximum dimension for Ca^2+^ bound form was 2.5 nm shorter than for the unbound form. G) Overlay of the Ca^2+^-bound P4H-TM SAXS data (red) with calculated SAXS data for monomeric (green) and dimeric (blue) crystal structure. H-I) Superposition of the dimeric P4H-TM crystal structure (black/gray) with the *ab initio* SAXS model of P4H-TM, H) without calcium (blue, χ^2^ 2.02) and I) with calcium (red, χ^2^ 1.69) generated with P2 symmetry.

To determine the difference in solution structure between Ca^2+^-bound and unbound P4H-TM, size-exclusion chromatography (SEC)-small-angle X-ray scattering (SAXS) was measured in the presence and absence of Ca^2+^. P4H-TM eluted as a single peak in both conditions (Fig. S5). Overall, the SAXS data obtained in the two conditions were not very different from each other (Fig. 7D), and there was no substantial difference in the degree of folding in the two conditions as evidenced by the Kratky plot (Fig. 7E). However, both the radius of gyration and the maximum dimension were larger for the sample without Ca^2+^ (Table 3, Fig. 7F). This result agrees with the models in Figs. 7A and 7C, where P4H-TM adopts a slightly extended conformation in the absence of calcium.

**Table 3.**
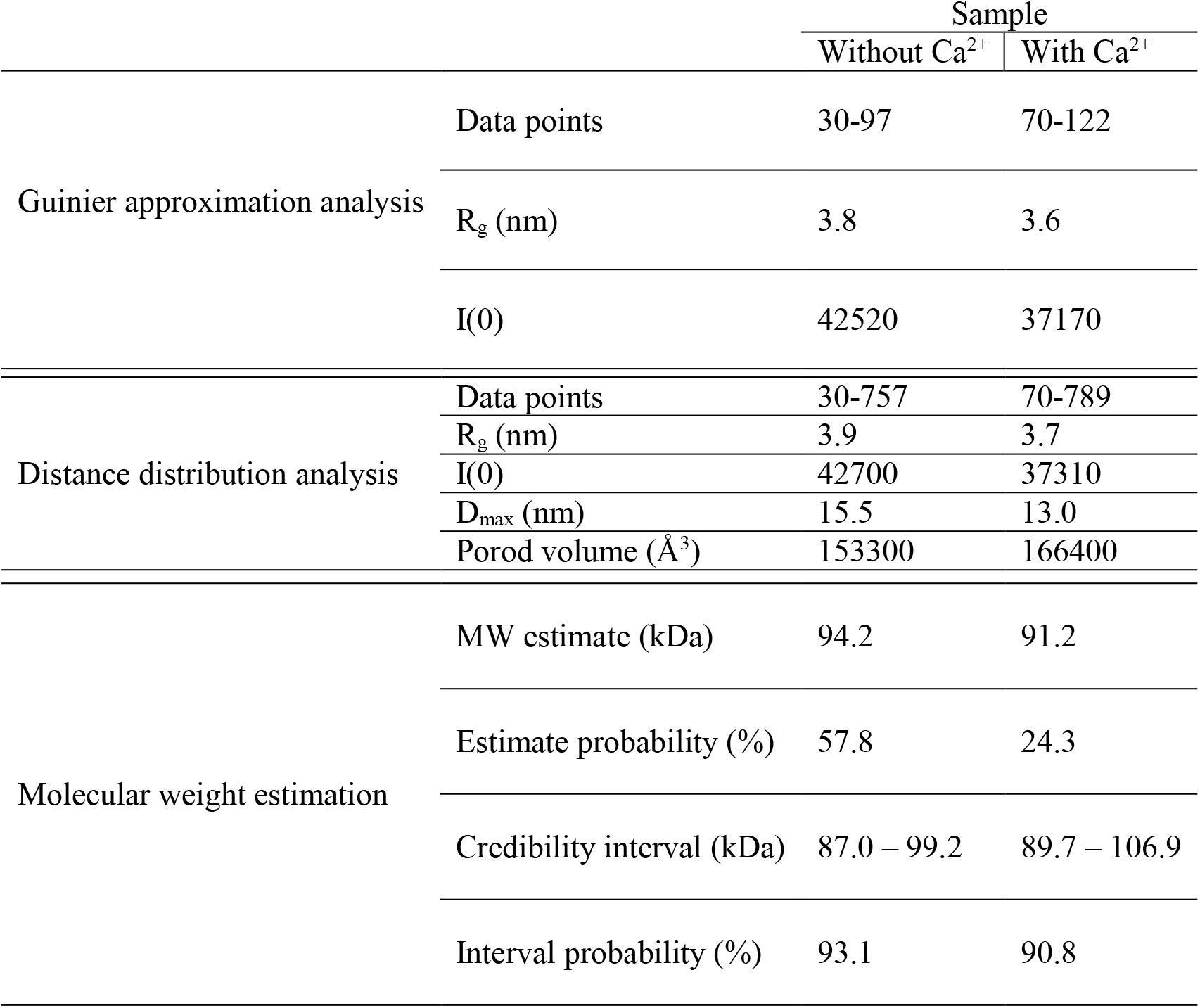
SAXS results of P4H-TM.

P4H-TM molecular weight calculated from the amino acid sequence is 48.3 kDa. The molecular weight estimates obtained from the SAXS data indicates that the protein eluted from the column as a dimer (Table 3). *Ab initio* modelling of the SAXS data with P2 symmetry produced extended envelopes with a central bulge (Fig. 7H and 7I, Fig. S6). The envelope of the protein in the absence of calcium is slightly longer corresponding to the larger D_max_. Superposition of the P4H-TM crystal structure with the SAXS envelopes suggests that the two molecules are positioned side-by-side at the central bulge and that the extended ends are formed by the N-terminus, the terminal α9 helix and the C-terminus. It seems likely that loss of calcium would cause a shift in the position of the dimerization interface resulting in a more elongated molecule. Such a shift could result from the movement of the EF-hand helices, as shown for example in Fig. 7C, but for a more detailed analysis, a higher resolution structure without calcium should be obtained.

To further study the oligomerization state, P4H-TM was analyzed by SEC-multi-angle light scattering (MALS) with and without Ca^2+^. Both samples contained a small amount of aggregated protein (Fig. S6). The molecular weight of the soluble protein fraction varied from 56-66 kDa without calcium to 60-70 kDa with calcium. These values are higher than the corresponding theoretical value of 48.3 kDa for the monomeric protein calculated from the amino acid sequence. In any case, the protein in the SEC-MALS assay eluted mainly in the monomeric form.

To obtain a third perspective on the oligomerization status, the P4H-TM crystal structure was analyzed using the PISA server. The results indicated that the complete structure of the dimer complex, including the metal and chloride ions, is thermodynamically stable. However, the results also stated that the interface between chains A and B is less hydrophobic than would be expected for a dimer interface and that it could be a crystal packing artefact.

### Mapping of the HIDEA variants to the crystal structure of P4H-TM

Five P4H-TM variants have been linked to a severe developmental HIDEA syndrome (Table 4) (21, 22). When modelled on the P4H-TM crystal structure solved here, three of the variants clearly destroy the function of the enzyme as they lead to nearly complete loss of the whole protein or large fragments of the catalytic domain. His161Pro introduces a proline residue in the place of a surface-facing histidine in the middle of α1 (Fig. 8). As prolines are unable to conform to α-helical geometry, this variant would produce a kink in α1 and probably disrupt its interaction with the DSBH major sheet next to it. A stop codon replacing Gln471 leads to the truncation of the protein by 32 residues and cuts off the latter half of α9 (Fig. 8). Although not visible in the crystal structure, the C-terminus of P4H-TM contains an ER retention signal, and its loss might cause P4H-TM to advance beyond ER in the secretory pathway.

**Table 4.** Prediction of the effect of the known pathological P4H-TM variants on protein function.

| Variant | Protein modification | Predicted outcome |
| --- | --- | --- |
| c.1073G>A | Arg296Ser + Val297-Arg358del | Loss of function |
| c.482A>C | His161Pro | Disruption of secondary structure |
| c.286dupC | Gln96Profs*29 | Loss of function |
| c.1594C>T | Gln471* | Loss of ER retention signal and disruption of secondary structure |
| c.949delG | Val317Phefs*30 | Loss of function |

**Fig. 8.**
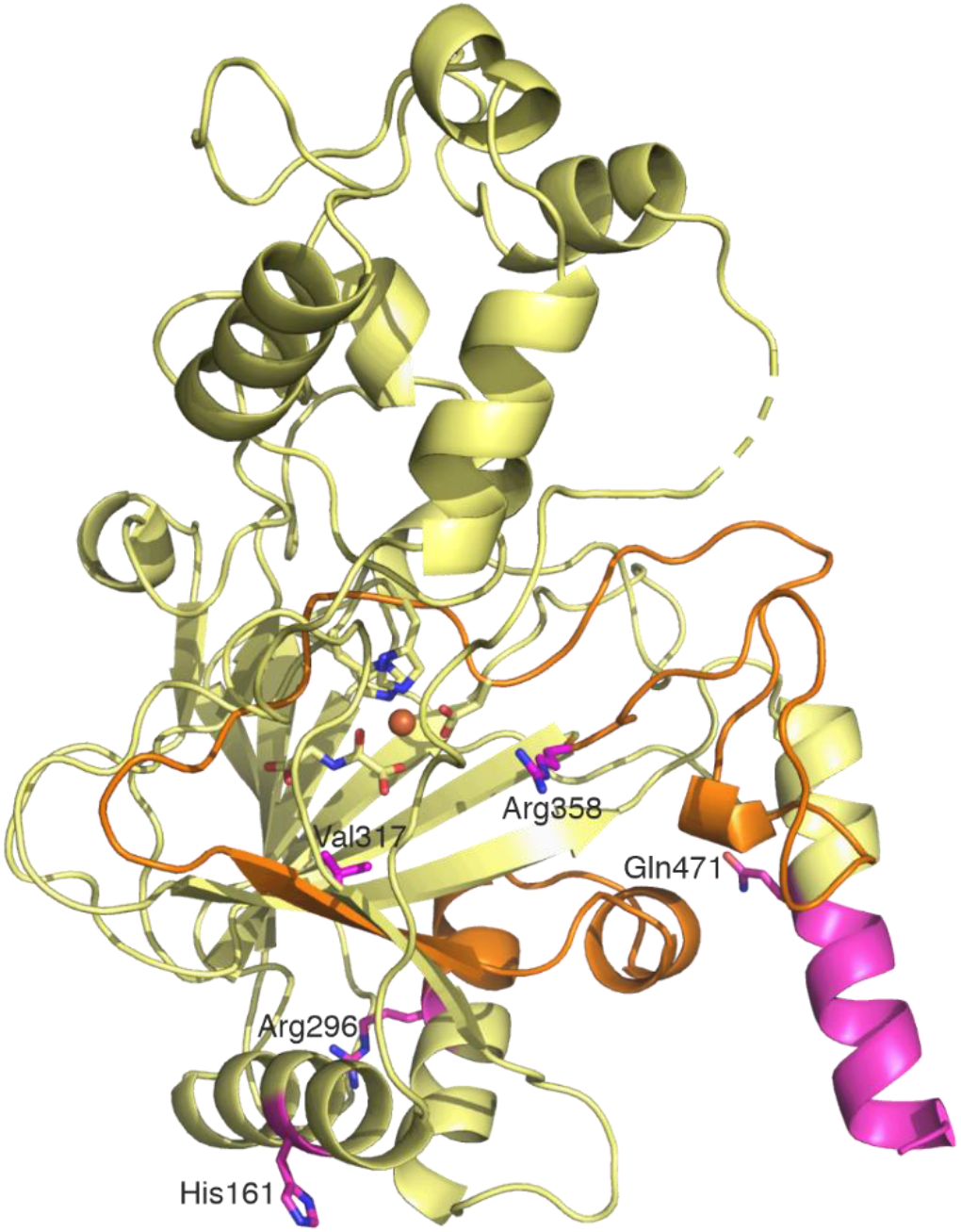
HIDEA variants presented in the P4H-TM crystal structure. Active site histidine and aspartate residues and N-oxalylglycine are shown in yellow. Specific HIDEA variant residues are presented in magenta. The region coded by exon 6 is shown in orange and the region truncated by the early stop codon replacing Gln471 is shown in magenta.

## Discussion

P4H-TM is a functionally enigmatic enzyme that is localized at the ER membrane (10, 11). It is composed of an N-terminal cytoplasmic tail, a membrane-anchoring transmembrane helix, and a unique combination of a Ca^2+^-binding EF-domain and a catalytic domain that is located within the ER lumen. The crystal structure reported here reveals the structure of the soluble part of P4H-TM without the cytoplasmic tail and the transmembrane helix. P4H-TM belongs to the 2OGDD family characterized by the DSBH structural fold. The helix-loop-helix structures of the EF-domain are inserted into the middle of the catalytic P4H domain but are structurally distinct from it. In between the catalytic domain and the EF-domain forms the substrate binding cavity of the enzyme. The cavity is located above the catalytic center which contains Fe^2+^ and the 2OG analog NOG. The 2OGDD protein family includes single-domain and large multi-domain proteins that are highly variable in their substrate specificity, but P4H-TM is the only enzyme in this family that has EF-hand motifs (23).

Previous SEC studies proposed that both the full-length P4H-TM and the construct used here (residues 88-502) are dimers with molecular weights around 105-120 kDa and around 85-90 kDa, respectively (11). The current study suggests that both monomeric and dimeric forms are possible. Two P4H-TM copies were found in the asymmetric unit in the P4H-TM crystals. PISA server analysis suggested that the packing found in the crystal is stable, but that the interaction interface is less hydrophobic than expected, implying that it could be a crystal packing artefact. The molecular weight from the SEC-MALS analysis was much closer to the weight of a monomeric than of a dimeric protein. In contrast, the solution structure determined with SEC-SAXS resembled the crystallographic dimer and the molecular weight estimated from the SEC-SAXS data indicated a dimer. However, the molecular weight of the SEC elution peak was not homogenous in the SEC-SAXS experiment. The input concentration of P4H-TM was higher in the SEC-SAXS experiment than in the SEC-MALS experiment, and the chromatography column was of a smaller volume in SEC-SAXS than in SEC-MALS, resulting in increased dilution in the latter. This suggests that the oligomerization could be concentration dependent. Another possible explanation for the discrepancy is that the SEC-SAXS sample was a mixture of monomers and dimers and the column resolution was not adequate to separate these two states. In the SEC-MALS experiments, a smaller peak was observed eluting before the main peak, possibly corresponding to the dimeric protein. The presence or absence of calcium made no difference to the oligomerization state. Based on these results, it is not possible to unequivocally conclude the oligomerization state of P4H-TM. Furthermore, there may be additional interaction sites within the transmembrane or cytoplasmic regions and P4H-TM may also exist as a transient dimer, as shown for some proteins such as the zebrafish SCP-2 thiolase (24). The monomer-dimer exchange could be important for the function of P4H-TM, since the SEC-SAXS model of the P4H-TM dimer suggested the enzyme to form an extended shape that would be attached to the membrane from both ends, and where both active sites of the dimer would be in a similar orientation adjacent to the ER membrane. Such a model highlights the fact that very little is currently known about the role and possible interaction partners of the cytoplasmic N-terminus of P4H-TM.

P4H-TM variants have been found to cause the HIDEA syndrome. Its symptoms include hypotonia, severe intellectual disability, epilepsy and eye abnormalities (21, 22). Some of the reported variants will clearly lead to P4H-TM loss of function as they result in the loss of large fragments of the enzyme, including the active site. For the variants His161Pro and Gln471* previous analysis indicated decreased protein solubility (22). The structural analysis revealed that His161 is located within the α1 helix in the P4H-TM structure. Introduction of a proline within the helix will produce a bend to the helix. α1 forms conserved hydrophobic interactions with the major sheet of the DSBH fold and the α7 helix, and a conserved salt bridge between helices α1 and α7. Therefore, disruption of α1 probably leads to the destabilization of the enzyme. The early stop codon replacing Gln471 removes a part of the C-terminal helix α9 and any subsequent residues that were not visible in the crystal structure. The sequence of the terminal helix and residues following Gln471 are not strongly conserved either in the P4H-TM homologs or orthologs suggesting the critical effect of this variant may be the loss of the ER retention signal at the C-terminus of the protein.

In addition to the 502-residue isoform reported initially (10, 11), some databases now include additional isoforms of P4H-TM that would be derived from alternative splicing. A 563-residue isoform 3, resulting from mis-splicing of exons 6 and 7, has been suggested to be the “canonical” isoform and has been used as the template for the antibody epitope for P4H-TM in Human Protein Atlas (http://www.proteinatlas.org/) (25). However, there is no published evidence of this isoform appears in protein form and there are no peptides listed in PeptideAtlas (26) that correspond to the 61 residues unique for this isoform. This 563-residue isoform was previously found not to be expressed in human fibroblasts or myoblasts (22). Further, we were unable here to express and purify this isoform using the insect cell expression system, and the interpretation of the P4H-TM crystal structure suggests that the 563-residue isoform would not preserve the conserved structural core of the enzyme. Together these data indicate that the 563-residue isoform is likely to be a splicing artefact that is not translated into a functional enzyme.

Two groups of P4Hs are found in animals. C-P4Hs hydroxylate specific prolines in procollagen chains and enable the formation of the stable triple-helical structure (2, 7). HIF-P4Hs hydroxylate two prolines in HIFα proteins leading to their proteasomal degradation (5, 8). P4H-TM has characteristics of both groups. It is localized to ER like C-P4Hs and its amino acid sequence is more similar to C-P4Hs than HIF-P4Hs (10, 11). On the other hand, it does not hydroxylate prolines in collagen or HIFα peptides but has some activity towards the ODDD of HIF1α (11), and it contributes to HIF1α degradation and regulation of EPO (19). However, it has not been thoroughly clarified what exactly is the role of P4H-TM in HIF regulation. In addition to animals, P4Hs are found in plants, algae, bacteria and viruses (27). A handful of residues in the P4H active site are conserved in all or nearly all of these enzymes. Although some of the conserved residues function to preserve the conserved structural fold, several of them are linked to the P4H catalytic activity. P4H-TM active site resembles the classical active site of 2OGDD family enzymes (28). It contains a divalent iron, central to the catalytic activity, coordinated by two histidines and an aspartate residue and the 2OG analog NOG. Although a precise enzymatic mechanism for any P4H has not been described, the similarities between the P4H-TM active site and the active sites of Cr-P4H and HIF-P4H-2 described here, extend beyond the iron and 2OG coordinating residues to the residues that interact with the substrate peptide and the proline to be hydroxylated. This confirms that P4H-TM is indeed a prolyl 4-hydroxylase, most likely with a peptide substrate.

P4H-TM structure revealed an extensive substrate binding cavity between the EF-domain and the catalytic domain. The cavity is bordered by loop structures extending from the DSBH core of the catalytic domain. The sequences of these loops are not conserved in the homologous P4Hs but are strongly conserved among the P4H-TM orthologs. Therefore, these loops are likely to participate in P4H-TM substrate binding or activity regulation. Cr-P4H and HIF-P4H-2 structures have been described both in the presence and absence of the peptide substrates and both enzymes have similar lid structures folding over the substrate peptide (16, 29–31). In this P4H-TM structure, the α1-α2 loop is partially disordered and positioned like a lid in an open conformation. It seems likely to be capable to form a lid structure over a substrate peptide when one is bound. In P4H-TM homolog structures the lid residues interact with the substrate peptide and in HIF-P4H-2 they are also known to contribute to the substrate specificity (29, 31). Interestingly, the residues of the lid structures are not conserved between P4H-TM and the other P4Hs. The analysis of the electrostatic surface near the P4H-TM active site, and its comparison with HIF-P4H-2, revealed that the P4H-TM active site contains abundant negative charge, while HIF-P4H-2 contains mainly positive charge. These data, together with the previous results showing that P4H-TM does not hydroxylate HIF-P4H substrate peptides, suggest that P4H-TM has evolved to bind a different substrate than HIF-P4H-2 (11).

The cell stores calcium in mitochondria and ER. The Ca^2+^ concentration within the ER lumen has been measured to be between 100 and 800 μM while concentrations as low as 1 μM have been reported in human cells during Ca^2+^ mobilization (32, 33). Ca^2+^ *K*_*d*_ for P4H-TM was measured here to be 23 μM, indicating that P4H-TM would be saturated with calcium in physiological situation, but that this saturation could be sensitive to changes, such as when calcium is temporarily released from the ER to the cytosol. EF-hand containing proteins often regulate enzyme activity in response to changes in cellular calcium concentration. However, the role of the EF-domain in P4H-TM is not currently fully understood. In a previous study, 5 mM CaCl_2_ was included in the activity assays where P4H-TM was found to be inactive towards HIF1α or collagen peptides and active towards the HIF1α ODDD (11). P4H-TM seems to be able to adopt both monomeric and dimeric forms in solution, but calcium had no effect on the oligomerization status. Some EF-hand motifs adopt an unstructured molten globule conformation in the absence of calcium and only form regular secondary structure when calcium is bound (13). However, SRCD measurements did not find any major shift in the secondary structure of P4H-TM when calcium was added. On the other hand, a near-UV CD measurement produced a major shift in the tryptophan-region at 285-305 nm, indicating a movement induced by the EF-hand calcium binding in the position of some of the tryptophans in the enzyme. This movement is likely to arise from the two tryptophans found in the α3-α4 loop between the two EF-hands that is expected to shift position when the enzyme adopts the calcium-bound conformation seen in the crystal structure. The α3-α4 tryptophans are located near the active site cavity and a conformation change in this region in response to calcium binding could indicate changes in the active site relevant to enzymatic activity. Some EF-hand motifs also bind magnesium (13). We did not see a shift in the near-UV CD signal when magnesium was added and we were unable to crystallize P4H-TM when calcium in the crystallization condition was replaced with magnesium, suggesting that P4H-TM does not bind magnesium. Since P4H-TM could not be crystallized in the absence of calcium, the effect of the loss of calcium on the P4H-TM structure was modelled based on the structure of the apo form of calmodulin. Three different models were generated where either 1) the initial α2 helix or 2) the subsequent α3 helix were fixed in place, or where 3) the α5 helix was modelled to extend the α6 helix. Two of these models led to an extended overall structure for P4H-TM where the EF-domain was seen to move away from the catalytic domain, causing widening of the active site cavity. Such a model would likely have an impact on the catalytic activity of P4H-TM. The SEC-SAXS results indicated that P4H-TM without calcium adopts a slightly extended conformation where both the radius of gyration and the maximum dimension are slightly larger compared to the calcium-bound P4H-TM. However, since P4H-TM in the SAXS data was found to be a dimer, it is not clear if the extended conformation is caused by the elongation of both monomers or the reorganization of the dimer interface.

In conclusion, the solved 3D structure of P4H-TM indicates that it shares the key structural elements of the known P4Hs confirming that it is a true P4H while possessing a unique property among the 2OGDDs having an EF-domain and a catalytic activity potentially regulated by Ca^2+^.

## Experimental procedures

### Cloning, expression and purification

The cloning of the P4H-TM construct has been described previously (11). Briefly, the construct contains 17 N-terminal residues (MLRRALLCLAVAALVRA) from the ER localization signal of protein disulfide isomerase that are cleaved upon import to the ER, six histidine residues and the residues 88-502 of human P4HTM within the pVL1392 expression vector. In order to improve the expression yield, this construct was subcloned into pFastBac Dual vector (Invitrogen) and Bac-to-Bac protocol was used with the EMBacY *Escherichia coli* strain (34) to generate P4H-TM bacmids. P4H-TM isoform 3 (MGC clone 3940241) was obtained from the Genome Biology Unit at the University of Helsinki, Finland. Isoform 3 DNA sequence was amplified with PCR, the amplified DNA and the P4H-TM isoform 1 plasmid were both digested with BlpI (NEB) and XbaI (NEB) restriction enzymes and the isoform 1 sequence was replaced with isoform 3. Bacmids were transfected to Sf9 cells using Baculofectin II transfection reagent (Oxford Expression Technologies). Resulting viruses were used to infect Sf21 expression cultures in insect-Xpress media (Lonza). Expression culture cells were harvested 48 hours after proliferation arrest, washed with PBS and frozen in −80°C.

Frozen cells were thawed and resuspended in lysis buffer containing 10 mM Tris-HCl pH 7.8, 0.1 M glycine, 0.1 M NaCl, 20 mM imidazole, 2 mM CaCl_2_, 20 μM FeSO_4_, 0.1% Triton X-100 and 1x protease inhibitor cocktail (Roche). The cell suspension was homogenized, the insoluble fraction was pelleted by centrifugation and the soluble fraction applied to a His-trap Ni^2+^-affinity column or a gravity-flow Ni-NTA column. The column was washed with 10 mM Tris-HCl pH 7.8, 0.1 M glycine, 0.1 M NaCl, 20 mM imidazole, 2 mM CaCl_2_, 20 μM FeSO_4_ and the bound proteins were eluted with similar buffer with 0.3 M imidazole. The eluted fractions were analyzed with SDS-PAGE and the P4H-TM containing fractions were pooled, concentrated and purified with size-exclusion chromatography (SEC) using 10 mM Tris-HCl pH 7.8, 0.1 M glycine, 0.1 M NaCl, 2 mM CaCl_2_, 20 μM FeSO_4_ as eluate. The protein samples used to analyze Ca^2+^ interaction were purified in the same way, but without CaCl_2_ or FeSO_4_ and 1 mM EDTA was added after elution from the Ni^2+^-affinity column.

### Multi-angle light scattering (MALS)

Molecular mass and sample quality of P4H-TM with and without Ca^2+^ were analyzed with a MiniDAWN MALS device (Wyatt Technology Corporation, Santa Barbara, USA) connected to a Shimadzu HPLC unit (Shimadzu Corporation, Kyoto, Japan) with a Superdex 200 Increase 10/300 GL SEC column (GE Healthcare Life Sciences) at constant 10°C temperature and equilibrated with the SEC buffer with and without 2 mM CaCl_2_. The sample concentration was 3.3 mg/ml. The RID-10A refractive index detector (Shimadzu Corporation) connected to the HPLC system was used as a concentration source for the calculations. ASTRA software (version 7.3.1.) (Wyatt Technology Corporation) was used to calculate the molecular weight and polydispersity of the samples.

### Circular dichroism (CD) spectroscopy

SRCD spectra were collected from 0.3 mg/ml samples at AU-CD beamline at ASTRID2 synchrotron source (ISA, Aarhus, Denmark) The samples were prepared to a buffer with 1 mM Tris-HCl pH 7.8, 10 mM NaCl and 10 mM glycine. 2 mM CaCl_2_ was added right before the measurement. The samples were equilibrated to room temperature and applied into 0.1 mm pathlength closed quartz cuvettes (Suprasil, Hellma Analytics). The spectra were recorded from 170 nm to 280 nm, at 25°C. Three repeat scans per measurement were recorded. The spectra were processed and the baselines were subtracted using CDToolX (35).

Near-UV CD spectra were collected using a Chirascan CD spectrometer (Applied Photophysics, Leatherhead, UK) between 250 and 350 nm at room temperature using a 1 cm path length quartz cuvette. The CD measurements were acquired every 1 nm with 1 s as an integration time and repeated three times with baseline correction. For the near-UV measurement, P4H-TM was diluted so that the absorbance at 280 nm was 1. The samples were measured so that the protein in SEC buffer without metal was measured first, after which CaCl_2_ or MgCl_2_ was added to the cuvette to 2 mM concentration, and the sample was measured again. The data were analyzed with Pro-Data Viewer (Applied Photophysics).

### Crystallization and data collection

P4H-TM crystals were grown using sitting drop vapor diffusion method. The drops (200 nl protein solution and 100 nl well solution) were made with the Mosquito nanodispenser (TTP Labtech) and imaged using the Formulatrix RI27 plate hotel at 4°C at the Structural biology core facility at Biocenter Oulu. The crystallization results were monitored using the in-house IceBear software (Daniel et al., manuscript in preparation). The protein concentration was 3 mg/ml, and the buffer was the same used for the SEC analyses including 2 mM CaCl_2_ and 20 μM FeSO_4_. The well solution was 0.1 M Tris-HCl pH 9, 22% tert-butanol and 1 mM N-oxalylglycine (NOG, Sigma). The crystals were soaked briefly in a solution containing 0.1 M Tris-HCl pH 9, 5% tert-butanol, 20% 2-methyl-2,4-pentanediol and 1 mM NOG, before flash-freezing in liquid nitrogen. Diffraction data were collected at the beamline P13 operated by EMBL Hamburg at the PETRA III storage ring (DESY, Hamburg, Germany) (36). Crystals suffered radiation damage during data collection and a minimal number of images (collected at the beginning of the exposure time) that produced a complete dataset were used in the final data processing calculations.

### Data processing and structure refinement

Data were processed in XDS (37). Molecular replacement was done with Phaser using a single molecule of Cr-P4H (PDB id 2jig), modified with Phenix.sculptor as search model (16, 38, 39). Two molecules were found in the asymmetric unit and the model was built initially using Phenix.autobuild followed by several cycles of manual building in COOT and structure refining using Phenix.refine (40–42). The resolution cutoff 2.25 Å was determined using paired refinement within PDB-redo server (43, 44). The structure was validated using Molprobity and PDB validation server (45, 46). The glycan conformations were validated using the pdb-care server (47). Data processing and refinement statistics are shown in Table 5.

**Table 5.** Data collection, data processing and structure refinement statistics. (Values in parenthesis are for the highest resolution shell)

|  |  |
| --- | --- |
| **Data collection statistics** |  |
| Beamline | P13 EMBL/DESY PETRA III |
| Detector | PILATUS 6M |
| Temperature (K) | 100 |
| Wavelength (Å) | 0.976 |
| Resolution range (Å) | 46.0 - 2.25 (2.33 - 2.25) |
| Space group | P 3<sub>1</sub> |
| Unit cell (Å) | 92.1 92.1 129.5 |
| (°) | 90 90 120 |
| Molecules per asymmetric unit | 2 |
| V<sub>m</sub> (Å<sup>3</sup>/Da) | 3.4 |
| Total reflections | 196716 (12872) |
| Unique reflections | 57041 (4799) |
| Multiplicity | 3.4 (2.7) |
| Completeness (%) | 98.0 (83.0) |
| Mean I/sigma(I) | 7.35 (0.40) |
| Wilson B-factor (Å<sup>2</sup>) | 60.8 |
| R<sub>p.i.m.</sub> | 0.057 (1.34) |
| CC<sub>1/2</sub> | 0.995 (0.18) |
| **Refinement statistics** |  |
| Resolution range (Å) | 50.3 – 2.25 |
| Reflections used in refinement | 57027 |
| Reflections used for R-free | 2008 |
| R-work | 0.182 |
| R-free | 0.221 |
| Number of non-hydrogen atoms | 6394 |
| Macromolecules | 5903 |
| Ligands | 356 |
| Waters | 135 |
| Protein residues | 726 |
| RMS(bonds) (Å) | 0.005 |
| RMS(angles) (°) | 0.7 |
| Ramachandran favored (%) | 97.9 |
| Ramachandran allowed (%) | 2.0 |
| Ramachandran outliers (%) | 0.1 |
| Rotamer outliers (%) | 0 |
| Clashscore | 5.4 |
| Average B-factor (Å<sup>2</sup>) | 87.8 |
| Proteins (Å<sup>2</sup>) | 86.6 |
| Ligands (Å<sup>2</sup>) | 116.0 |
| Waters (Å<sup>2</sup>) | 63.5 |
| Number of TLS groups | 8 |

### Structure analysis

Structure figures were generated with PyMOL (Schrödinger, LLC) and UCSF Chimera (48). APBS plugin for PyMOL was used to generate the electrostatic surfaces (49). The secondary-structure matching algorithm was used in COOT to align the homologous structures of Cr-P4H (PDB id: 2jig and 3gze), HIF-P4H-1 (PDB id: 5v1b), HIF-P4H-2 (PDB id: 3hqr and 2g19) and Ba-P4H (PDB id: 5hv4) to P4H-TM (16, 29, 30, 50–53). P4H-TM calcium-free morph structures were based on the structure of rat apo calmodulin (PDB id: 1qx5) N-terminal EF-hand pair (54). Different helices of the calmodulin EF-hand motifs were aligned with the corresponding helices in P4H-TM EF-domain using the LSQ algorithm in COOT. P4H-TM EF-domain was then morphed to resemble the calcium-free calmodulin using UCSF Chimera. The packing of the two P4H-TM molecules in the crystal structure was analyzed using the PISA server (55). The structure based sequence alignment was done using PSI-Search and Clustal-Omega (56, 57). The sequence editing and annotations were done using Genedoc (58).

### Isothermal titration calorimetry (ITC)

P4H-TM and Ca^2+^ interaction was studied using ITC at the Proteomics and protein analysis core facility at Biocenter Oulu. Ca^2+^-free P4H-TM was exchanged with SEC to a buffer containing 10 mM Tris-HCl pH 7.8, 0.1 M glycine and 0.1 M NaCl. CaCl_2_ was dissolved in the same buffer, diluted to 4 mM and injected to 120 μM P4H-TM at 25°C using ITC200 instrument (MicroCal, Malvern, UK). The binding was analyzed with MicroCal Origin using ‘one set of sites’ binding model.

### Small-angle X-ray scattering (SAXS)

SAXS measurements were done at the P12 beamline at PETRA III in Hamburg (59). P4H-TM purified in the absence of calcium was passed through Superdex 200 Increase 5/150 column with 0.2 ml/min flow rate at room temperature. Scattering was measured from the eluted buffer and protein sections. Identical samples were applied to the system in two buffers which both contained 10 mM Tris-HCl pH 7.8, 0.1 M NaCl, 0.1 M glycine and 1% (w/v) glycerol, and one of which also contained 2 mM CaCl_2_. The protein and buffer frames were selected for processing using CHROMIXS (60). Buffer frames were averaged, and the averaged buffer intensity was subtracted from the individual protein frames. The subtracted protein frames were then scaled and averaged. The data were processed and the Bayesian inference molecular weight estimates were obtained using PRIMUS (61, 62). 20 *ab initio* models were generated for both datasets using GASBOR (63). The models were averaged using DAMAVER and the most typical model was selected (64). Experimental scattering curves were compared with curves calculated from the P4H-TM structure monomer and dimer with CRYSOL (65). The processing software were part of the ATSAS package version 3 (66).

## Data availability

P4H-TM crystal structure has been submitted to Protein Data Bank with the identification number 6tp5.

## Acknowledgements

We thank Eeva Lehtimäki and Essi Kivilahti for expert technical assistance. The Biocenter Oulu structural biology, proteomics and protein analysis, and sequencing center facilities and their expertise is gratefully acknowledged. The synchrotron MX data and SAXS data were collected at beamlines P13 and P12, respectively, operated by EMBL Hamburg at the PETRA III storage ring (DESY, Hamburg, Germany). We would like to thank Johanna Hakanpää and Karen Manalastas for the assistance in using the beamlines. The use of ASTRID2 AU-CD beamline (ISA, Aarhus, Denmark) was supported by the project CALIPSOplus under the Grant Agreement 730872 from the EU Framework Programme for Research and Innovation HORIZON 2020. The P4H-TM isoform 3 clone 3940241 was obtained from the MGC Library; Genome Biology Unit supported by HiLIFE and the Faculty of Medicine, University of Helsinki, and Biocenter Finland.

## Abbreviations

2OG: 2-oxoglutarate
2OGDD: 2-oxoglutarate-dependent dioxygenase
CD/SRCD: circular dichroism/synchrotron radiation circular dichroism
C-P4H: collagen prolyl 4-hydroxylase
Cr-P4H: *C. reinhardtii* prolyl 4-hydroxylase
DSBH: double-stranded β-helix
EPO: erythropoietin
ER: endoplasmic reticulum
HIF: hypoxia-inducible factor
HIF-P4H: HIF prolyl 4-hydroxylase
ITC: isothermal titration calorimetry
MALS: multi-angle static light scattering
NOG: N-oxalylglycine
ODDD: oxygen-dependent degradation domain
P4H-TM: transmembrane prolyl 4-hydroxylase
P4H: prolyl 4-hydroxylase
SAXS: small-angle X-ray scattering
SEC: size-exclusion chromatography

## References

1. McDonough, M. A., Loenarz, C., Chowdhury, R., Clifton, I. J., and Schofield, C. J. (2010) Structural studies on human 2-oxoglutarate dependent oxygenases. Curr. Opin. Struct. Biol. 20, 659–672.

2. Myllyharju, J. (2003) Prolyl 4-hydroxylases, the key enzymes of collagen biosynthesis. Matrix Biol. J. Int. Soc. Matrix Biol. 22, 15–24.

3. Myllyharju, J., and Kivirikko, K. I. (2004) Collagens, modifying enzymes and their mutations in humans, flies and worms. Trends Genet. 20, 33–43.

4. Kaelin, W. G., and Ratcliffe, P. J. (2008) Oxygen sensing by metazoans: the central role of the HIF hydroxylase pathway. Mol. Cell. 30, 393–402.

5. Myllyharju, J., and Koivunen, P. (2013) Hypoxia-inducible factor prolyl 4-hydroxylases: common and specific roles. Biol. Chem. 394, 435–448.

6. Koski, M. K., Anantharajan, J., Kursula, P., Dhavala, P., Murthy, A. V., Bergmann, U., Myllyharju, J., and Wierenga, R. K. (2017) Assembly of the elongated collagen prolyl 4-hydroxylase α2β2 heterotetramer around a central α2 dimer. Biochem. J. 474, 751–769.

7. Rappu, P., Salo, A. M., Myllyharju, J., and Heino, J. (2019) Role of prolyl hydroxylation in the molecular interactions of collagens. Essays Biochem. 63, 325–335.

8. Hirsilä, M., Koivunen, P., Günzler, V., Kivirikko, K. I., and Myllyharju, J. (2003) Characterization of the human prolyl 4-hydroxylases that modify the hypoxia-inducible factor. J. Biol. Chem. 278, 30772–30780

9. Koivunen, P., Hirsilä, M., Kivirikko, K. I., and Myllyharju, J. (2006) The length of peptide substrates has a marked effect on hydroxylation by the hypoxia-inducible factor prolyl 4-hydroxylases. J. Biol. Chem. 281, 28712–28720.

10. Oehme, F., Ellinghaus, P., Kolkhof, P., Smith, T. J., Ramakrishnan, S., Hütter, J., Schramm, M., and Flamme, I. (2002) Overexpression of PH-4, a novel putative proline 4-hydroxylase, modulates activity of hypoxia-inducible transcription factors. Biochem. Biophys. Res. Commun. 296, 343–349.

11. Koivunen, P., Tiainen, P., Hyvärinen, J., Williams, K. E., Sormunen, R., Klaus, S. J., Kivirikko, K. I., and Myllyharju, J. (2007) An endoplasmic reticulum transmembrane prolyl 4-hydroxylase is induced by hypoxia and acts on hypoxia-inducible factor alpha. J. Biol. Chem. 282, 30544–30552.

12. Kretsinger, R. H., and Nockolds, C. E. (1973) Carp Muscle Calcium-binding Protein II. STRUCTURE DETERMINATION AND GENERAL DESCRIPTION. J. Biol. Chem. 248, 3313–3326.

13. Gifford, J. L., Walsh, M. P., and Vogel, H. J. (2007) Structures and metal-ion-binding properties of the Ca^2+^-binding helix-loop-helix EF-hand motifs. Biochem. J. 405, 199–221.

14. Kawasaki, H., and Kretsinger, R. H. (2017) Structural and functional diversity of EF-hand proteins: Evolutionary perspectives. Protein Sci. 26, 1898–1920.

15. Taylor, M. S. (2001) Characterization and comparative analysis of the EGLN gene family. Gene. 275, 125–132.

16. Koski, M. K., Hieta, R., Böllner, C., Kivirikko, K. I., Myllyharju, J., and Wierenga, R. K. (2007) The active site of an algal prolyl 4-hydroxylase has a large structural plasticity. J. Biol. Chem. 282, 37112–37123

17. Hyvärinen, J., Parikka, M., Sormunen, R., Rämet, M., Tryggvason, K., Kivirikko, K. I., Myllyharju, J., and Koivunen, P. (2010) Deficiency of a transmembrane prolyl 4-hydroxylase in the zebrafish leads to basement membrane defects and compromised kidney function. J. Biol. Chem. 285, 42023–42032.

18. Leinonen, H., Rossi, M., Salo, A. M., Tiainen, P., Hyvärinen, J., Pitkänen, M., Sormunen, R., Miinalainen, I., Zhang, C., Soininen, R., Kivirikko, K. I., Koskelainen, A., Tanila, H., Myllyharju, J., and Koivunen, P. (2016) Lack of P4H-TM in mice results in age-related retinal and renal alterations. Hum. Mol. Genet. 25, 3810–3823.

19. Laitala, A., Aro, E., Walkinshaw, G., Mäki, J. M., Rossi, M., Heikkilä, M., Savolainen, E.-R., Arend, M., Kivirikko, K. I., Koivunen, P., and Myllyharju, J. (2012) Transmembrane prolyl 4-hydroxylase is a fourth prolyl 4-hydroxylase regulating EPO production and erythropoiesis. Blood. 120, 3336–3344.

20. Leinonen, H., Koivisto, H., Lipponen, H.-R., Matilainen, A., Salo, A. M., Dimova, E. Y., Hämäläinen, E., Stavén, S., Miettinen, P., Myllyharju, J., Koivunen, P., and Tanila, H. (2019) Null mutation in P4h-tm leads to decreased fear and anxiety and increased social behavior in mice. Neuropharmacology. 153, 63–72.

21. Kaasinen, E., Rahikkala, E., Koivunen, P., Miettinen, S., Wamelink, M. M. C., Aavikko, M., Palin, K., Myllyharju, J., Moilanen, J. S., Pajunen, L., Karhu, A., and Aaltonen, L. A. (2014) Clinical characterization, genetic mapping and whole-genome sequence analysis of a novel autosomal recessive intellectual disability syndrome. Eur. J. Med. Genet. 57, 543–551.

22. Rahikkala, E., Myllykoski, M., Hinttala, R., Vieira, P., Nayebzadeh, N., Weiss, S., Plomp, A. S., Bittner, R. E., Kurki, M. I., Kuismin, O., Lewis, A. M., Väisänen, M.-L., Kokkonen, H., Westermann, J., Bernert, G., Tuominen, H., Palotie, A., Aaltonen, L., Yang, Y., Potocki, L., Moilanen, J., van Koningsbruggen, S., Wang, X., Schmidt, W. M., Koivunen, P., and Uusimaa, J. (2019) Biallelic loss-of-function P4HTM gene variants cause hypotonia, hypoventilation, intellectual disability, dysautonomia, epilepsy, and eye abnormalities (HIDEA syndrome). Genet. Med. 21, 2355–2363.

23. Aik, W., McDonough, M. A., Thalhammer, A., Chowdhury, R., and Schofield, C. J. (2012) Role of the jelly-roll fold in substrate binding by 2-oxoglutarate oxygenases. Curr. Opin. Struct. Biol. 22, 691–700.

24. Kiema, T.-R., Thapa, C. J., Laitaoja, M., Schmitz, W., Maksimainen, M. M., Fukao, T., Rouvinen, J., Jänis, J., and Wierenga, R. K. (2019) The peroxisomal zebrafish SCP2-thiolase (type-1) is a weak transient dimer as revealed by crystal structures and native mass spectrometry. Biochem. J. 476, 307–332.

25. Thul, P. J., Åkesson, L., Wiking, M., Mahdessian, D., Geladaki, A., Ait Blal, H., Alm, T., Asplund, A., Björk, L., Breckels, L. M., Bäckström, A., Danielsson, F., Fagerberg, L., Fall, J., Gatto, L., Gnann, C., Hober, S., Hjelmare, M., Johansson, F., Lee, S., Lindskog, C., Mulder, J., Mulvey, C. M., Nilsson, P., Oksvold, P., Rockberg, J., Schutten, R., Schwenk, J. M., Sivertsson, Å., Sjöstedt, E., Skogs, M., Stadler, C., Sullivan, D. P., Tegel, H., Winsnes, C., Zhang, C., Zwahlen, M., Mardinoglu, A., Pontén, F., von Feilitzen, K., Lilley, K. S., Uhlén, M., and Lundberg, E. (2017) A subcellular map of the human proteome. Science. 10.1126/science.aal3321

26. Deutsch, E. W., Sun, Z., Campbell, D., Kusebauch, U., Chu, C. S., Mendoza, L., Shteynberg, D., Omenn, G. S., and Moritz, R. L. (2015) State of the Human Proteome in 2014/2015 As Viewed through PeptideAtlas: Enhancing Accuracy and Coverage through the AtlasProphet. J. Proteome Res. 14, 3461–3473.

27. Gorres, K. L., and Raines, R. T. (2010) Prolyl 4-hydroxylase. Crit. Rev. Biochem. Mol. Biol. 45, 106–124.

28. Hausinger, R. P. (2004) FeII/alpha-ketoglutarate-dependent hydroxylases and related enzymes. Crit. Rev. Biochem. Mol. Biol. 39, 21–68.

29. Koski, M. K., Hieta, R., Hirsilä, M., Rönkä, A., Myllyharju, J., and Wierenga, R. K. (2009) The crystal structure of an algal prolyl 4-hydroxylase complexed with a proline-rich peptide reveals a novel buried tripeptide binding motif. J. Biol. Chem. 284, 25290–25301.

30. McDonough, M. A., Li, V., Flashman, E., Chowdhury, R., Mohr, C., Liénard, B. M. R., Zondlo, J., Oldham, N. J., Clifton, I. J., Lewis, J., McNeill, L. A., Kurzeja, R. J. M., Hewitson, K. S., Yang, E., Jordan, S., Syed, R. S., and Schofield, C. J. (2006) Cellular oxygen sensing: Crystal structure of hypoxia-inducible factor prolyl hydroxylase (PHD2). Proc. Natl. Acad. Sci. U. S. A. 103, 9814–9819.

31. Chowdhury, R., Leung, I. K. H., Tian, Y.-M., Abboud, M. I., Ge, W., Domene, C., Cantrelle, F.-X., Landrieu, I., Hardy, A. P., Pugh, C. W., Ratcliffe, P. J., Claridge, T. D. W., and Schofield, C. J. (2016) Structural basis for oxygen degradation domain selectivity of the HIF prolyl hydroxylases. Nat. Commun. 7, 12673

32. Burdakov, D., Petersen, O. H., and Verkhratsky, A. (2005) Intraluminal calcium as a primary regulator of endoplasmic reticulum function. Cell Calcium. 38, 303–310.

33. Miyawaki, A., Llopis, J., Heim, R., McCaffery, J. M., Adams, J. A., Ikura, M., and Tsien, R. Y. (1997) Fluorescent indicators for Ca^2+^ based on green fluorescent proteins and calmodulin. Nature. 388, 882–887.

34. Bieniossek, C., Richmond, T. J., and Berger, I. (2008) MultiBac: Multigene Baculovirus-Based Eukaryotic Protein Complex Production. Curr. Protoc. Protein Sci. 51, 5.20.1–5.20.26

35. Miles, A. J., and Wallace, B. A. (2018) CDtoolX, a downloadable software package for processing and analyses of circular dichroism spectroscopic data. Protein Sci. 27, 1717–1722.

36. Cianci, M., Bourenkov, G., Pompidor, G., Karpics, I., Kallio, J., Bento, I., Roessle, M., Cipriani, F., Fiedler, S., and Schneider, T. R. (2017) P13, the EMBL macromolecular crystallography beamline at the low-emittance PETRA III ring for high- and low-energy phasing with variable beam focusing. J. Synchrotron Radiat. 24, 323–332.

37. Kabsch, W. (2010) XDS. Acta Crystallogr. D Biol. Crystallogr. 66, 125–132.

38. Bunkóczi, G., and Read, R. J. (2011) Improvement of molecular-replacement models with Sculptor. Acta Crystallogr. D Biol. Crystallogr. 67, 303–312.

39. McCoy, A. J., Grosse-Kunstleve, R. W., Adams, P. D., Winn, M. D., Storoni, L. C., and Read, R. J. (2007) Phaser crystallographic software. J. Appl. Crystallogr. 40, 658–674.

40. Terwilliger, T. C., Grosse-Kunstleve, R. W., Afonine, P. V., Moriarty, N. W., Zwart, P. H., Hung, L. W., Read, R. J., and Adams, P. D. (2008) Iterative model building, structure refinement and density modification with the PHENIX AutoBuild wizard. Acta Crystallogr. D Biol. Crystallogr. 64, 61–69.

41. Emsley, P., Lohkamp, B., Scott, W. G., and Cowtan, K. (2010) Features and development of Coot. Acta Crystallogr. D Biol. Crystallogr. 66, 486–501.

42. Afonine, P. V., Grosse-Kunstleve, R. W., Echols, N., Headd, J. J., Moriarty, N. W., Mustyakimov, M., Terwilliger, T. C., Urzhumtsev, A., Zwart, P. H., and Adams, P. D. (2012) Towards automated crystallographic structure refinement with phenix.refine. Acta Crystallogr. D Biol. Crystallogr. 68, 352–367

43. Karplus, P. A., and Diederichs, K. (2012) Linking Crystallographic Model and Data Quality. Science. 336, 1030–1033.

44. Joosten, R. P., Long, F., Murshudov, G. N., and Perrakis, A. (2014) The PDB_REDO server for macromolecular structure model optimization. IUCrJ. 1, 213–220.

45. Chen, V. B., Arendall, W. B., Headd, J. J., Keedy, D. A., Immormino, R. M., Kapral, G. J., Murray, L. W., Richardson, J. S., and Richardson, D. C. (2010) MolProbity: all-atom structure validation for macromolecular crystallography. Acta Crystallogr. D Biol. Crystallogr. 66, 12–21.

46. Gore, S., Sanz García, E., Hendrickx, P. M. S., Gutmanas, A., Westbrook, J. D., Yang, H., Feng, Z., Baskaran, K., Berrisford, J. M., Hudson, B. P., Ikegawa, Y., Kobayashi, N., Lawson, C. L., Mading, S., Mak, L., Mukhopadhyay, A., Oldfield, T. J., Patwardhan, A., Peisach, E., Sahni, G., Sekharan, M. R., Sen, S., Shao, C., Smart, O. S., Ulrich, E. L., Yamashita, R., Quesada, M., Young, J. Y., Nakamura, H., Markley, J. L., Berman, H. M., Burley, S. K., Velankar, S., and Kleywegt, G. J. (2017) Validation of Structures in the Protein Data Bank. Structure. 25, 1916–1927.

47. Lütteke, T., and von der Lieth, C.-W. (2004) pdb-care (PDB carbohydrate residue check): a program to support annotation of complex carbohydrate structures in PDB files. BMC Bioinformatics. 5, 69

48. Pettersen, E. F., Goddard, T. D., Huang, C. C., Couch, G. S., Greenblatt, D. M., Meng, E. C., and Ferrin, T. E. (2004) UCSF Chimera--a visualization system for exploratory research and analysis. J. Comput. Chem. 25, 1605–1612.

49. Baker, N. A., Sept, D., Joseph, S., Holst, M. J., and McCammon, J. A. (2001) Electrostatics of nanosystems: application to microtubules and the ribosome. Proc. Natl. Acad. Sci. U. S. A. 98, 10037–10041

50. Krissinel, E., and Henrick, K. (2004) Secondary-structure matching (SSM), a new tool for fast protein structure alignment in three dimensions. Acta Crystallogr. D Biol. Crystallogr. 60, 2256–2268.

51. Chowdhury, R., McDonough, M. A., Mecinović, J., Loenarz, C., Flashman, E., Hewitson, K. S., Domene, C., and Schofield, C. J. (2009) Structural basis for binding of hypoxia-inducible factor to the oxygen-sensing prolyl hydroxylases. Structure. 17, 981–989.

52. Schnicker, N. J., and Dey, M. (2016) Structural analysis of cofactor binding for a prolyl 4-hydroxylase from the pathogenic bacterium Bacillus anthracis. Acta Crystallogr. Sect. Struct. Biol. 72, 675–681.

53. Ahmed, S., Ayscough, A., Barker, G. R., Canning, H. E., Davenport, R., Downham, R., Harrison, D., Jenkins, K., Kinsella, N., Livermore, D. G., Wright, S., Ivetac, A. D., Skene, R., Wilkens, S. J., Webster, N. A., and Hendrick, A. G. (2017) 1,2,4-Triazolo-[1,5-a]pyridine HIF Prolylhydroxylase Domain-1 (PHD-1) Inhibitors With a Novel Monodentate Binding Interaction. J. Med. Chem. 60, 5663–5672.

54. Schumacher, M. A., Crum, M., and Miller, M. C. (2004) Crystal structures of apocalmodulin and an apocalmodulin/SK potassium channel gating domain complex. Structure. 12, 849–860.

55. Krissinel, E., and Henrick, K. (2007) Inference of macromolecular assemblies from crystalline state. J. Mol. Biol. 372, 774–797.

56. Li, W., McWilliam, H., Goujon, M., Cowley, A., Lopez, R., and Pearson, W. R. (2012) PSI-Search: iterative HOE-reduced profile SSEARCH searching. Bioinformatics. 28, 1650–1651.

57. Sievers, F., Wilm, A., Dineen, D., Gibson, T. J., Karplus, K., Li, W., Lopez, R., McWilliam, H., Remmert, M., Söding, J., Thompson, J. D., and Higgins, D. G. (2011) Fast, scalable generation of high-quality protein multiple sequence alignments using Clustal Omega. Mol. Syst. Biol. 7, 539

58. Nicholas, K. B., Nicholas, H. B. J., and Deerfield, D. W. (1997) GeneDoc: Analysis and Visualization of Genetic Variation. EMBnet.news. 4, 1–4.

59. Blanchet, C. E., Spilotros, A., Schwemmer, F., Graewert, M. A., Kikhney, A., Jeffries, C. M., Franke, D., Mark, D., Zengerle, R., Cipriani, F., Fiedler, S., Roessle, M., and Svergun, D. I. (2015) Versatile sample environments and automation for biological solution X-ray scattering experiments at the P12 beamline (PETRA III, DESY). J. Appl. Crystallogr. 48, 431–443.

60. Panjkovich, A., and Svergun, D. I. (2018) CHROMIXS: automatic and interactive analysis of chromatography-coupled small-angle X-ray scattering data. Bioinformatics. 34, 1944–1946.

61. Konarev, P. V., Volkov, V. V., Sokolova, A. V., Koch, M. H. J., and Svergun, D. I. (2003) *PRIMUS* : a Windows PC-based system for small-angle scattering data analysis. J. Appl. Crystallogr. 36, 1277–1282.

62. Hajizadeh, N. R., Franke, D., Jeffries, C. M., and Svergun, D. I. (2018) Consensus Bayesian assessment of protein molecular mass from solution X-ray scattering data. Sci. Rep. 8, 7204

63. Svergun, D. I., Petoukhov, M. V., and Koch, M. H. J. (2001) Determination of domain structure of proteins from x-ray solution scattering. Biophys. J. 80, 2946–2953.

64. Volkov, V. V., and Svergun, D. I. (2003) Uniqueness of *ab initio* shape determination in small-angle scattering. J. Appl. Crystallogr. 36, 860–864.

65. Svergun, D., Barberato, C., and Koch, M. H. J. (1995) *CRYSOL* – a Program to Evaluate X-ray Solution Scattering of Biological Macromolecules from Atomic Coordinates. J. Appl. Crystallogr. 28, 768–773.

66. Franke, D., Petoukhov, M. V., Konarev, P. V., Panjkovich, A., Tuukkanen, A., Mertens, H. D. T., Kikhney, A. G., Hajizadeh, N. R., Franklin, J. M., Jeffries, C. M., and Svergun, D. I. (2017) ATSAS 2.8: a comprehensive data analysis suite for small-angle scattering from macromolecular solutions. J. Appl. Crystallogr. 50, 1212–1225.

